# Atomistic Molecular Dynamics Simulations of Trioleoylglycerol – Phospholipid Membrane Systems

**DOI:** 10.1101/2020.01.25.918136

**Authors:** Mirza Ahmed Hammad, Hafiza Minal Akram, Muhammad Sohail Raza

## Abstract

Adiposomes are phospholipid coated triacylglyceride particles that serve as structural models of the fat storage compartments of cells, known as lipid droplets (LDs); however, unlike LDs, they do not carry proteins. There is a deficit of available methods and experimental data regarding the internal packing of the adiposomes, and computer simulations offer a promising way to pinpoint the molecular arrangements within these structures. However, in the absence of a triacylglycerol-specific atomic forcefield, thus far, all adiposome/LD simulations have been performed with the coarse grained/united atom forcefields. Yet it is desirable to model the phospholipid/triacylglycerol interface with atomic resolution. In the present study, we first prepared a 2-monooleoylglycerol (MOG) forcefield which was then used to build a trioleoylglycerol (TOG) forcefield by the modular approach of the AMBER software suite. TOG bilayer membrane (2L) systems were modelled from two different initial conformations; TOG3 and TOG2:1. The simulations revealed that TOG2:1 is the most populated conformation in TOG membranes, irrespective of the starting conformation. Some other parameter optimizations were performed for TOG membranes based on which adiposome mimicking tetralayer membrane system (4L) was prepared with a TOG bilayer at core surrounded by two DOPC leaflets. The 4L membranes were stable throughout the simulations, however it was observed that a small amount of cations and water diffused from surface to the TOG core of the membrane. Based on these results a TAG-packing model was also developed. It is expected that the availability of MOG forcefield will equip future studies with a framework for molecular dynamics simulations of adiposomes/LDs.

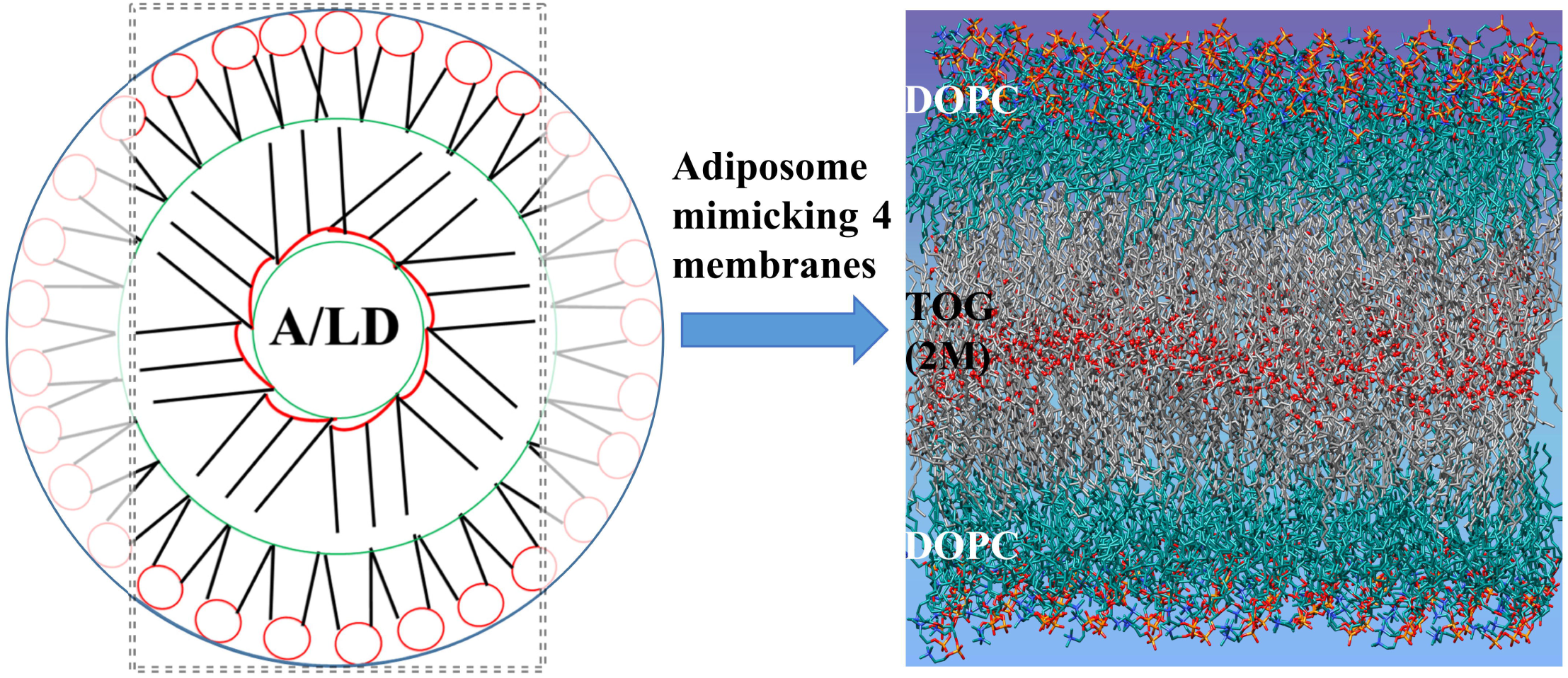

## Introduction

Adiposomes are created with a neutral lipid core, containing mostly triacylglycerols (TAGs), covered by a phospholipid monolayer to provide a simplified structural model of natural lipid droplets (LDs), the fat storage compartments of cells (1). Natural LDs serve as a platform for the function of a class of specialized proteins; adiposomes are usually constructed without these proteins (2). The study of LDs, and thus that of LD-mimetic adiposomes (artificial lipid droplets), is important due to their function in lipid homeostasis and - related - involvement in different diseases (3, 4); however adiposomes could also serve as an efficient pharmaceutical drug carrier system (5, 6). A key question is the basis of specific LD-binding of proteins to adiposomes/LDs (A/LD) that is directly related to the structure of the phospholipid-TAG interface. However, current state-of-the art experimental methods are unable to provide information about the molecular packing with high enough resolution. Therefore only computer simulation techniques provide the means to predict the molecular structure of the LD (7) and provide essential details (Supplementary Figure S1).

Few simulation studies have been performed to find out structural and dynamic properties of LD/apolipoprotein particles. These studies has been proved useful by providing results comparable to experimental outcomes. For example, modeling of high density lipoprotein particles (HDL) with cholesteryl esters (CEs) showed solubility behavior in accordance with experimental values (8). It was also observed that CEs mostly present in extended conformation and can move from core to surface to interact with water. Whilst in comparison with mixed TAG-CE HDL particles it was observed that TAGs has 25-50% higher ability than CEs to interact with aqueous medium (9–11). During oil/water modeling to represent lipid emulsion, authors showed that TAGs interacted with phospholipid monolayers resulting in exposure of water to TAGs (12). During lipid emulsion single and mixed phospholipid monolayers were more ordered than respective bilayers due to TAG-phospholipid tail-tail intercalations (13). In an attempt to find answers to TAG aggregation inside ER bilayer for LD formation (14), it was observed that TAGs were able to self-aggregate spontaneously in isotropic, mobile, disordered manner. It was suggested that such blisters would eventually leave ER when reaching specific size.

Many other aspects of A/LDs were also been studied to understand dynamic behavior of lipids i.e. from vesiculation of adiposomes to cholesterol - TAG interactions and interaction of proteins with LDs in some cases. It was shown that lipids tend to form vesicles when provided certain conditions (15). The vesicles can be in round shape as A/LDs or in cylindrical form if provided with different parameters. Triolein based A/LD simulations demonstrated the interaction of triolein with cholesterol molecules (16). It was observed that most cholesterol molecules tend to make their own clusters or micelles with triolein within A/LD. The same group demonstrated that some CEs also showed interaction with phospholipid monolayer of A/LD (17). Beside protein-less A/LD simulation, LD simulations with proteins have also been performed to determine LD-protein interactions. For example, CIDEA protein was shown to interact with LD monolayer through its amphipathic helix that possibly could facilitate TAG movement across LDs during LD-LD fusion (18). LD-protein interactions were also studied by another independent study where simulations were performed for LD and amphipathic helix of CCT α protein (19). It was shown that amphipathic helix of CCTα could interact with lipid monolayers due to LD packing defects. The caveolin-1 (cav1) protein was also shown to be involved in increasing curvature of LDs (20). Interestingly, it was shown that cav1 protein could interact with TAGs at neutral lipid-monolayer interface during LD formation and thins the bilayer at edges reducing energy barrier to pinch droplet off.

There are two dominant conformations of TAGs in an A/LD: the TAG3 type conformation where all lipid tails align in the same direction, and the TAG2:1 type conformation where two lipid tails point in one direction and the third to the opposite. There is experimental evidence that TAGs can form lamellar phases with the individual molecules in TOG2:1 conformation (21, 22). In the sole computational study on the conformations of TAGs in an A/LD it was noted that different TAG conformations were stable during the simulation (23). According to available crystal structures, TAGs form lamellar phases where the most probable conformation is TAG2:1 (24–26). It is unclear, however, whether this conformation of TAG is stable in LDs under provided constraints of the system and the higher (physiological) temperatures. When seeking answers to such questions, atomic level studies of TAGs are indispensable due to their ability to provide details with higher resolution and utile with actual experiments. With added point, till now no atomic level study of LD/apolipoprotein particles has been performed. One of the impediments of atomic level modelling studies was the absence of an atomic level forcefield for TAG. In current study, we, i) constructed such a forcefield for Trioleoylglycerol (TOG), a type of TAG with three oleic tails attached to glycerol ii) modelled two dominant TOG conformations iii) optimized different parameters for TOGs to simulate them with phospholipid monolayer iv) performed atomic level simulations of an adiposome-mimetic 4-layer (4L) system where the TOG bilayer is sandwiched between DOPC monolayers.

## Computational Methods

Initial simulations were run with pmemd.cuda of AMBER14 (www.ambermd.org) on Tesla C2050 GPU based computer system, provided by HPC of Institute of Biophysics CAS, Beijing, China. Additional calculations on newer GPUs (nVidia 1050 Ti and nVidia 1080 Ti) were performed with AMBER18 at the National Laboratory for Macrobiomolecular Research, Institute of Biophysics, CAS, Beijing, China.

Advanced Model Building with Energy Refinement (AMBER) package uses modular approach to build lipid molecules so that it has separate forcefields for oleoic (OL) tails and the glycerophospholipid headgroup, such as phosphatidyl choline (PC) (27). According to AMBER’s modular approach, DOPC can be constructed by combining PC headgroup with two OL tails without prior bond formation. The bond between the headgroup and two lipid tails would be automatically formed during the parametrization process. TOG and DOPC molecules show many basic structural similarities, so AMBER can be used to build TOG molecule based on DOPC structure (Figure 1A, 1B). The glycerol backbone, containing three carbon atoms C1, C2, and C3 with oxygen atoms O11, O21, and O31 is the same in DOPC and TOG. For DOPC, O11 and O21 oxygen atoms form esters with C11 and C21 of the OL tails, whereas O31 connects to P31 of the PC headgroup. For making a neutral lipid forcefield, from now on termed MOG, a triester was formed with two formic acid moieties and an oleoyl chain. According to crystal structures and available literature, the alkyl chain at C2 position is important for neutral lipids (28–30). Hence, to calculate the forcefield the OL tail was attached to C21 position while the former C11 and C31 would receive OL tail according to AMBER’s built-in modular approach after the MOG forcefield was calculated. This approach abates the need of calculating forcefield for the complete TOG molecule. The forcefield parameters for MOG were calculated through quantum mechanical (QM) calculations. RED webserver was employed for this purpose (31). Gaussian was used for QM calculations with standard method (32, 33). Charge neutrality of MOG molecule was obtained by adding protons where needed. Charge-refitting was performed to optimize atomic charges according to the geometry of MOG.

**Figure 1:**
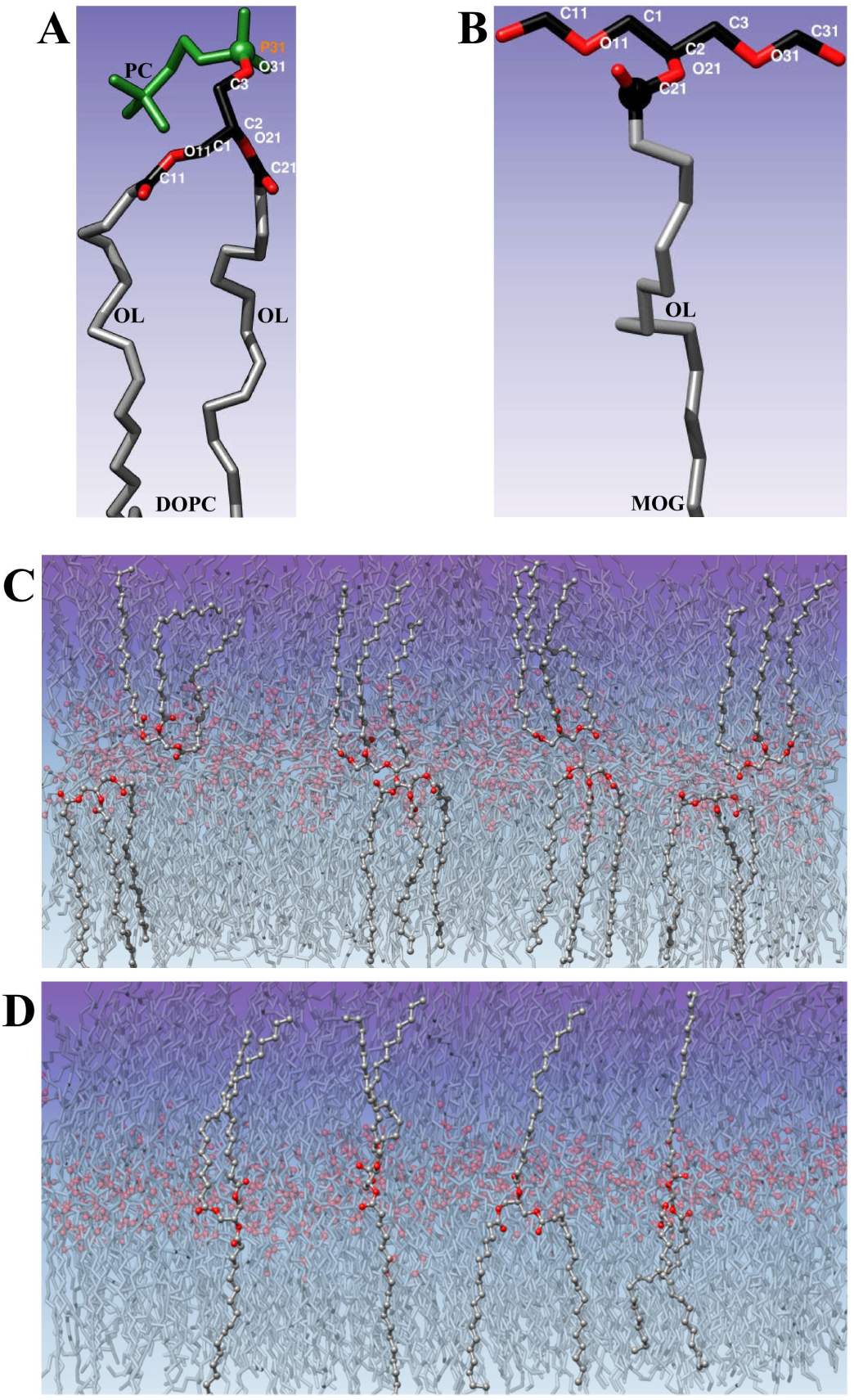
Preparation of MOG forcefield and TOG membranes. **Figure 1A/B:** Structures of DOPC and MOG molecules are shown. The substructure except OL lipid tails of both molecules is similar, similar carbon atoms are colored black and similar oxygen atoms are colored red. The P31 of DOPC headgroup (represented as color: green and atom type: sphere) was replaced with C21 (represented as color: black and atom type: sphere) along with OL lipid tail. Atomic level MOG forcefield parameters were calculated through QM simulations employing Gaussian tool from R.E.D webserver. Two lipid tails of DOPC were transferred to build complete TOG molecule through AMBER’s modular approach. **Figure 1C/D:** TOG membranes in two different conformations are shown. TOG3 2M system (1C) was prepared through memgen and AMBER’s built-in module AMBAT. AMBAT module was edited to contain TOG3 molecule with same parameters as for DOPC. It was not possible to prepare TOG2:1 2M system (1D) with memgen and AMBAT due to layer type membrane structure. Manual TOG2:1 membrane preparation was performed through UCSF Chimera tool utilizing movement mouse mode and transform coordinate commands from utilities section.

TOG molecule was built by attaching two OL tails from already available AMBER lipid17 repository to MOG by removing hydrogens at corresponding positions. When prepared, MOG had zero charge but due to removal of hydrogens and addition of OL tails, TOG molecule as a whole got (−0.14) charge. Based on available literature, two conformations for TOG molecule were created: i) TOG3 with 3 lipid tails in one direction and ii) TOG2:1 with two lipid tails in one direction and one lipid tail in opposite direction. The TOG3 type membranes were prepared through memgen (34) and AMBAT (35) tools (Figure 1C). The membrane structure for TOG2:1 conformation should be in the form of intercalating layers; memgen and AMBAT tools do not provide such functionality. Therefore TOG 2:1 type membranes were prepared semi-manually by UCSF Chimera (36, 37) (Figure 1D). The TOG membranes were prepared considering the same area per lipid/volume as a DOPC membrane; for TOG3 this yielded membranes of 144 molecules per layer, for TOG 2:1 91 molecules per layer due to the different conformation. The MOG library, TOG2:1, and TOG3 PDB formatted files for preparing membranes are provided in supplementary information (Supplementary File S2).

TOG bilayer systems (from here on, 2L) were optimized with AMBAT tool of AMBER with TOG atom information. For the adiposome-mimetic (4L) system, DOPC bilayer in water environment was prepared through AMBAT tool and TOG membrane was then placed between the leaflets. TOG 2L simulations were conducted in sequences where after each 100 ns run on pmemd.cuda (38), a 1 ns simulation was run using the sander.MPI module of AMBER (39). For 4L system, after each 20 ns simulation on pmemd.cuda, a 0.5 ns run was performed on sander.MPI. This strategy was needed to minimize the probability of local energy minima traps because with every new simulation the random seed was provided for the calculations (40, 41). Both TOG3 2L system and TOG2:1 2L system were first energy minimized and then simulated each at 273K, 277K, and 298K. To achieve stable systems at these temperatures, slow heating was applied so that systems were heated from 0K to 100K, 100K to 200K, and 200K to final temperatures (Supplementary Table S3). Production molecular dynamics (PMD) was performed for approximately 200 ns for each membrane system at each temperature under either constant pressure (NPT) or constant volume (NVT) ensemble with periodic boundary conditions. As in a system all TOGs were similar, isotropic NPT conditions were applied using Monte Carlo barostat. Same strategy was applied for all successive 2L systems with PMD of approximately 300 ns. For 4L system, water solvent along with 1 Cl^−^ and 36 Na^+^ ions were added to system and after minimization slow heating was performed up to 273K to avoid membrane deformations due to sudden temperature change. The heating was first applied to the water molecules while putting restraints on all other atoms. Following that the restraints were slowly relaxed modelling a process where heat flows from water to the core of 4L system. Initially, 100 ns simulation was performed for both systems but system under NVT ensemble showed continuous increase in RMSD. Therefore, the simulation for NVT ensemble was extended up to 500 ns. All the analyses were performed in cpptraj module of AMBER (42, 43). Graphs were made with xmgrace (44) and Microsoft Excel.

## Results

### Optimization of TOG bilayers

TOG 2L membranes in TOG3 and TOG2:1 conformations were simulated after preparation of complete TOG atomic level forcefield. For initial simulations, the TOG membranes were prepared considering the same area per lipid/volume as of 12×12 DOPC membrane. It was observed that TOG molecules exhibited random movements. RMSD graphs are shown in Figure 2A. The systems showed higher RMSD initially but gradually stabilized as observed through convergence of RMSD. The simulation at 298K showed higher random movements with higher RMSD while at 273K and 277K the movements were relatively less random and relatively lower RMSD. It was observed that at 273K the first RMSD peak was at less than 10 Å for both systems and it was above 10 Å at 277K. For 298K the first peak was above 20 Å for TOG3 2L system and about at 10 Å for TOG2:1 2L system. It suggests higher structural changes at the start of PMD that evolves towards a more stable assembly with time.

**Figure 2:**
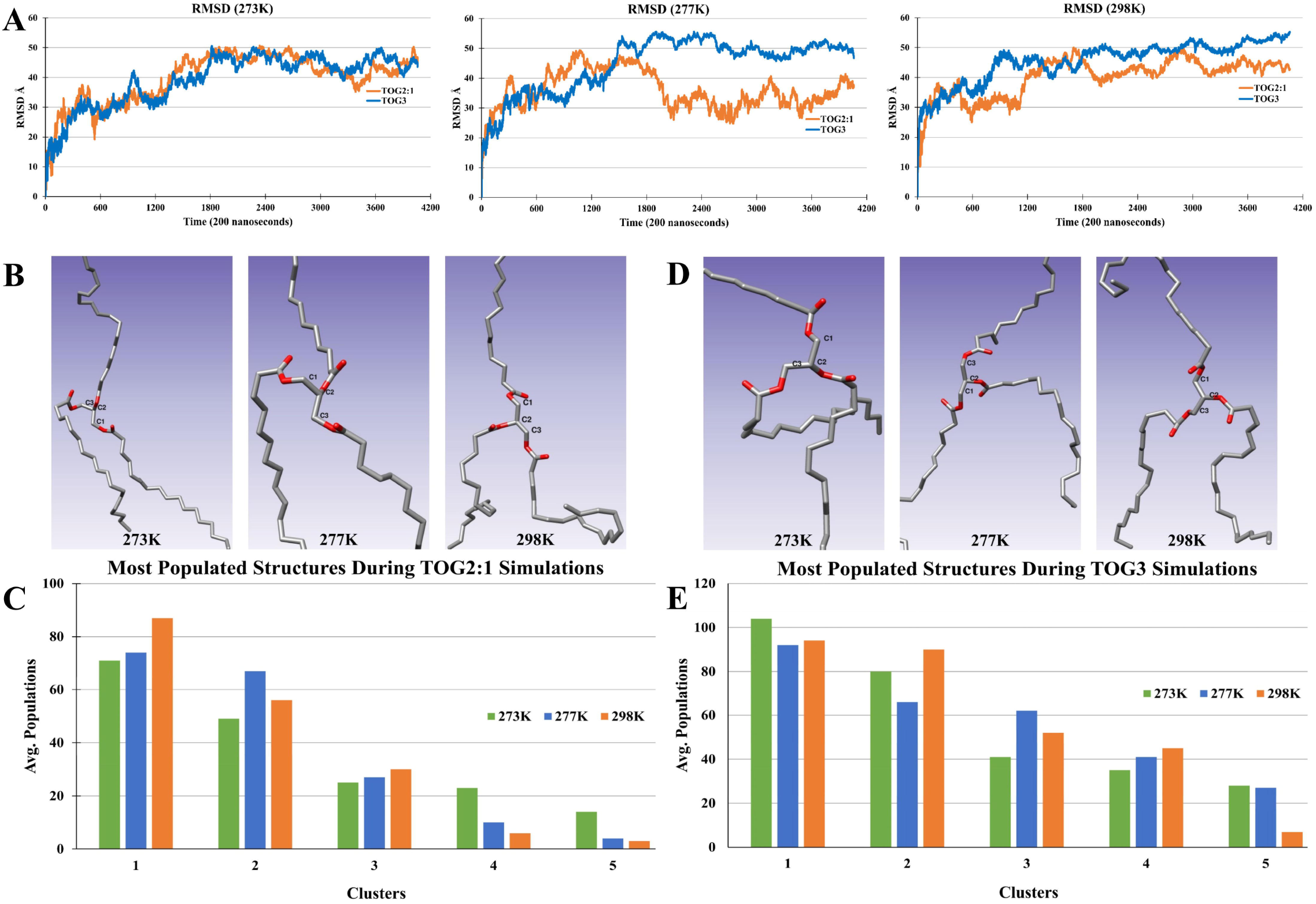
Comparison of TOG2:1 and TOG3 conformations based 2M systems. **Figure 2A:** RMSD graphs of simulations performed at three temperatures, 273K, 277K, and 298K. RMSD for only PMD steps of 200 ns is shown. The first peak of RMSD for TOG2:1 and TOG3 was at less than 10Å at 273K while RMSD peaks were higher at 277K and 298K. It can be observed that RMSD of TOG2:1 2M was lowest than TOG3 2M at all temperatures. Clustering analysis was performed to find out most populated structures for individual TOG molecules at different temperatures. kmeans clustering algorithm was employed for clustering. **Figure 2B/C:** The most populated structures for TOG2:1 2M system (2B) along with clusters (2C) are shown. Only structures of first cluster at each temperature is shown. **Figure 2D/E:** The most populated structures for TOG3 2M (2D) and clustering results (2E) are shown. Only structures of first cluster at each temperature is shown. Interestingly, it was observed that the most populated structure for TOG3 2M system was in TOG2:1 conformation (also please refer to Supplementary video S4).

The structural changes in individual TOG molecules from TOG3 and TOG2:1 were analyzed at each temperature. It was observed that after a few nanoseconds some TOG3 molecules changed their conformation to TOG2:1 and remained in this conformation to the end of simulation (Supplementary Video S4). Cluster analysis was performed to check whether this effect also prevails in all simulation systems at different temperatures. Cluster analysis performed with kmeans algorithm for all individual TOGs in each systems yielded a total of 6 trajectories; 3 trajectories for TOG2:1 2L system at 273K, 277K, and 298K based on 182 highest ranked TOG structures and 3 trajectories for TOG3 2L system at 273K, 277K, and 298K containing 288 highest ranked TOG structures. kmeans clustering was performed again over the trajectories of the highest ranked structures. The clustering analysis confirmed that TOGs in TOG2:1 2L system retained their conformation over time (Figure 2B, 2C) while TOGs in TOG3 2L system changed their conformation to TOG2:1 increasingly at each temperature steps (Figure 2D, 2E). These results suggest that TOG2:1 is the more stable conformation for TOGs and therefore all following simulations were based on TOG2:1 2L system.

The TOG molecules have high carbon ratio resulting in higher collisions at higher temperature. These high collisions had made it difficult to observe the behavior of individual molecules due to fast and random movements. Even at 273K the TOG molecules showed fast and random movements but were relatively lower than at 277K and 298K simulation. For this reason all consecutive simulations were carried out at 273K.

### Parameter Optimization

The previous TOG 2L systems were prepared with the same area per lipid/volume as for 12×12 DOPCs but it was observed that the box size or volume of system decreased significantly during heating step. It may affect TOG and DOPC interactions when simulated together. To remove any artefacts resulting from modelling TOG spacing on DOPC, calculations were performed with varied TOG-TOG distances. Keeping the same area, 3 systems of TOGs (in TOG2:1 conformation) were prepared with average 2-dimensioanl (2D) distances of 12 Å (system I), 6 Å (system II), and 3 Å (system III). Details are shown in Table 1. After minimization of each system, heating and PMD was performed. The simulations were performed with the same parameters for comparability (Supplementary S3). It was observed that during the heating step, system I with average 2D distance of 12 Å decreased the volume dramatically and system II with average 2D distance of 6 Å between each TOG also showed moderate decrease in volume. However system III with average 2D distance of 3 Å between each TOG increased its volume (Figure 3A). System I showed a continuous decrease in the radius of gyration while system II first showed decrease and then increase resulting an almost unchanged radius of gyration at the end of the simulation. System III showed an increase in radius of gyration (Figure 3B). The RMSD analysis showed that system II had relatively lower and stable RMSD values then system I and system III (Figure 3C). It appears from RMSD graph that system II stabilized at the very beginning of the PMD. Thus these results suggested that system II with average TOG-TOG 2D distance of 6 Å would provide the best representation of TOG membranes for the A/LD simulations in combination with DOPC.

**Table 1:**

| Experiment 1: Conformation based |  |  |  |  |
| --- | --- | --- | --- | --- |
| Components | Number | Details | Number of Atoms | Comments |
| TOG | 288 | 144 per layer | 48096 | TOG3 conformation |
| Charge of System | -39.4 | Neutralizing plasma | 0 |  |
| Total | 288 |  | 48096 |  |
| TOG | 182 | 91 per layer | 30394 | TOG2:1 conformation |
| Charge of System | -24.9 | Neutralizing plasma | 0 |  |
| Total | 182 |  | 30394 |  |
| Experiment 2: Number of TOGs in given volume |  |  |  |  |
| TOG | 182 | 91 per layer | 30394 | System I (TOG-TOG average distance of 12Å) |
| Charge of System | -24.9 | Neutralizing plasma | 0 |  |
| Total | 182 |  | 30394 |  |
| TOG | 254 | 127 per layer | 42418 | System II (TOG-TOG average distance of 6Å) |
| Charge of System | -34.8 | Neutralizing plasma | 0 |  |
| Total | 254 |  | 42418 |  |
| TOG | 398 | 199 per layer | 66466 | System III (TOG-TOG average distance of 3Å) |
| Charge of System | -54.5 | Neutralizing plasma | 0 |  |
| Total | 398 |  | 66466 |  |
| Experiment 3: Ensemble based |  |  |  |  |
| TOG | 254 | 127 per layer | 42418 | NPT and NVT Ensembles |
| Charge of System | -34.8 | Neutralizing plasma | 0 |  |
| Total | 254 |  | 42418 |  |
| Experiment 4: Adiposome mimicking 4L system |  |  |  |  |
| DOPC | 288 | 144 per layer | 39744 | NPT and NVT Ensembles |
| TOG | 254 | 127 per layer | 42418 |  |
| Water | 13037 | 6518 per layer | 39111 |  |
| Ions | 37 | 1 Cl- 36 Na+ | 37 |  |
| Total | 13616 |  | 121310 |  |

**Figure 3:**
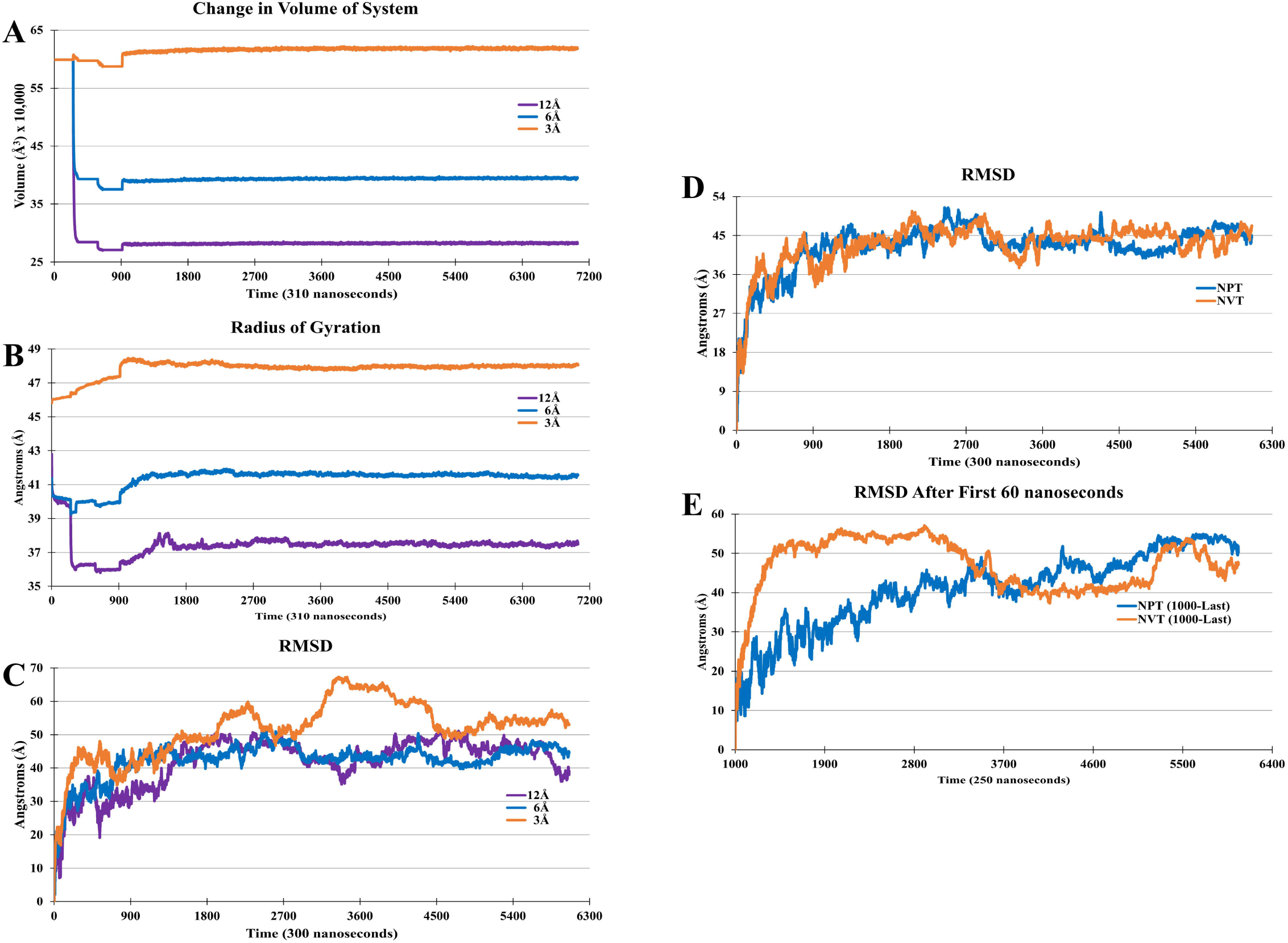
Parameter optimization for TOG 2M system including number of TOGs and ensemble. **Figure 3A/B/C:** The TOGs were arranged in a volume similar to 12×12 DOPC membrane with different TOG-TOG distances. Total three systems were prepared with average 2D distance between each TOG molecule i) 12Å (system I), ii) 6Å (system II), and iii) 3Å (system III). It was observed that during heating step, volume of system (3A) with average 2D distance 12Å decreased drastically while volume for system with average 2D distance 3Å increased. The volume for system with average distance 6Å also decreased but in middle range. Along volume, the radius of gyration (3B) graphs for average concentration of TOGs demonstrated that system with average 2D distance of 6Å showed better representation of TOG 2M system. The RMSD for PMD steps (3C) demonstrated that system with average 2D distance 6Å got stabilized earlier than other systems. **Figure 3D/E:** TOGs were simulated under NPT and NVT ensembles. The RMSD graphs (3D) demonstrated that both systems were converged with almost same RMSD values. But visually systems showed different behaviors for first 1000 frames (60 ns) (please refer to Supplementary video S5). The RMSD after first 1000 frames (3E) showed that system under NVT ensemble had higher RMSD earlier but also got stabilized earlier. Whereas system under NPT ensemble was not converged and constantly showed increasing RMSD.

The effect of ensemble parameters was investigated by comparing the stability of TOG membranes after heating under NPT and NVT (Table 1 and Supplementary S3). The RMSD for PMD steps showed that both systems have converged from their initial conformation showing very similar behavior (Figure 3D). A detailed look however revealed that TOG membranes under NVT ensemble showed different structural changes than under NPT ensemble (Supplementary Video S5). It was observed that under NVT ensemble, TOGs were arranged initially in a curved plane so that some TOGs from the edges were moving upward while TOGs in middle remained at the bottom. This behavior lasted for ~60 ns and after ~60 ns the TOGs started to move randomly diminishing the curved structure. It could indicate that there is a stable curved membrane conformation for TOGs that could not develop possibly due to the relatively small number of molecules or possibly small simulation box. For NPT ensemble, the TOG movements were continuously random without any structure. The RMSD after first 60 ns show that TOGs under NVT ensemble became stabilized after the diminishing of the curved structure while TOGs under NPT ensemble showed constantly increasing RMSD (Figure 3E). It suggests that for simulation of TOGs, NVT ensemble is better suited and may provide useful information about vesiculation of A/LDs.

### Simulation of 4L System

The TOG bilayers obtained from previous results were used to prepare tetralayer system where TOGs served as core layers and DOPCs as outer layers. The resulting 4L system mimicking adiposome cross section is shown in Figure 4A. After minimization of 4L system and heating from 0 K to 100 K, the membrane thickness was 79 Å including the 43.3 Å thick TOG membrane (Figure 4A). As TOG 2L membranes were simulated in both NVT and NPT ensembles, to have a clear understanding of ensemble effect, two simulations were performed for 4L system under NPT and NVT ensembles (Table 1 and Supplementary S3). Initially, 100 ns simulation was performed for both systems but system under NVT ensemble showed continuous increase in RMSD. So, simulation for NVT ensemble was extended up to 500 ns but still RMSD did not converged (data not shown).

**Figure 4:**
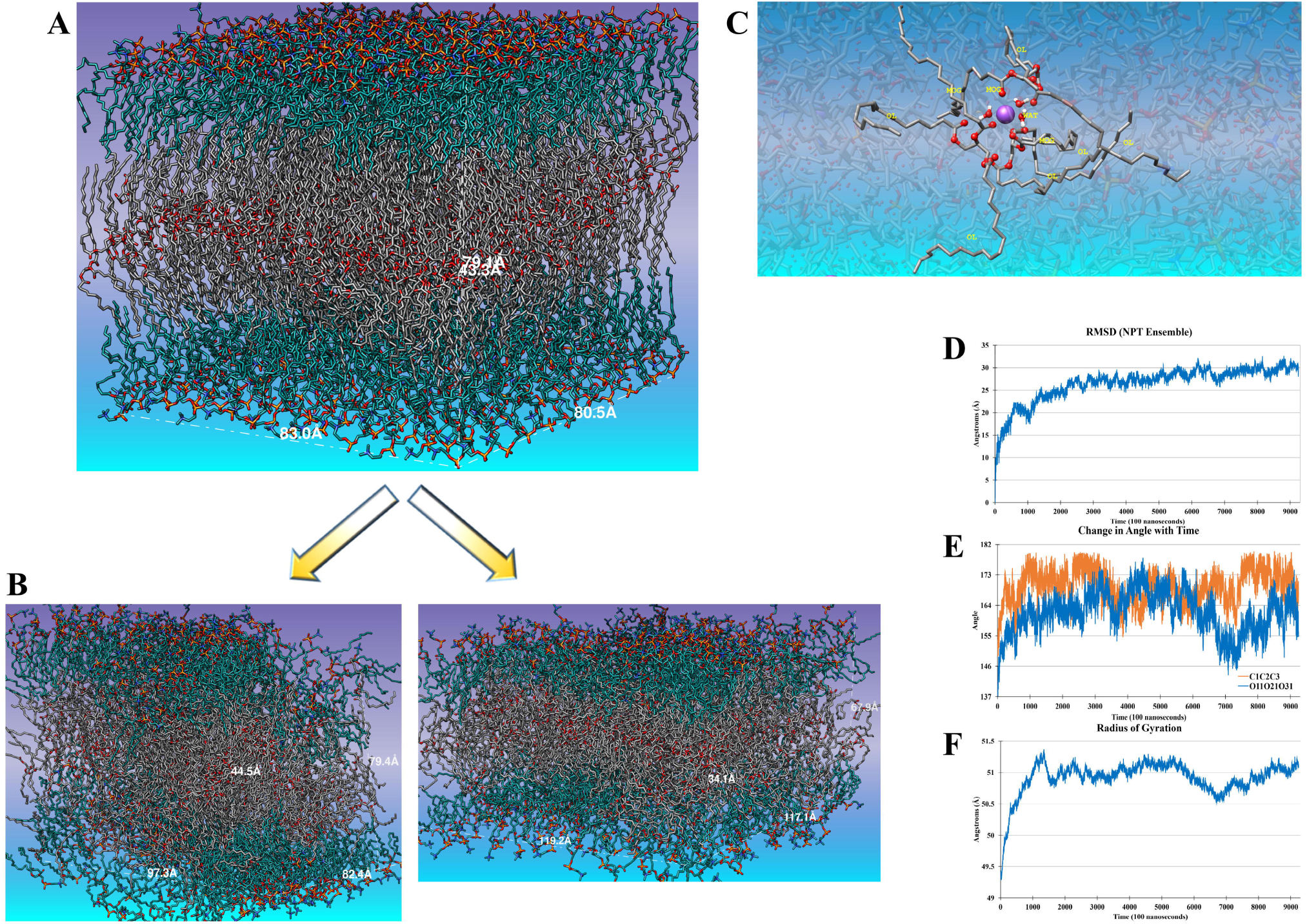
Simulation of adiposome mimicking 4 membrane system. **Figure 4A:** Adiposome mimicking 4M system after heating up to 100K is shown. The length and width of system is 80Å and 83Å respectively. The height is 79.1Å including 43.3Å TOG layer in middle. **Figure 4B:** 4M system was simulated under NVT ensembles (right) and NPT (left). The system under NVT ensemble was simulated for 500 ns but still showed constant increase in RMSD (data not shown). While system under NPT ensemble showed high TOG-DOPC interactions which resulted in changed box dimensions reducing its height and increasing length and width. **Figure 4C:** During simulation run, it was observed that some Na^+^ ions moved inside core of 4M system along with water. This behavior was higher in NVT ensemble and lower in NPT ensemble. At most three TOGs were observed to cover one Na^+^ ion (figure data is from NPT ensemble). **Figure 4D/E/F:** RMSD graph (4D) for 4M system under NPT ensemble showed that RMSD was stable and almost converged while reaching 100 ns simulation time. The change in angles of TOG core atoms (4E) and radius of gyrations of TOG molecules (4F) demonstrated that after few nanoseconds the TOGs may have some changes which forced the system to change its box dimensions.

It was observed that the membrane swelled within the available volume in NVT ensemble (Figure 4B, left hand), therefore the membrane in NVT ensemble was more fluid at 273K temperature. Partial intercalation was observed between the oleoyl moieties of TOG and DOPC lipids, hence the properties of the 4L system are necessarily influenced by these alkyl-alkyl interactions. On other hand, in the NPT ensemble (Figure 4B, right hand) the volume of system *decreased* drastically during the heating steps and PMD. TOG and DOPC molecules exhibited high degree of tail-tail intercalation. Several TOG molecules diffused from the membrane core to the DOPC layer. The simulations also showed an intrusion of ions and water molecules to the core of the TOG layers. This was observed in both NPT and NVT ensemble simulations. It was observed that each Na^+^ ion was surrounded by up to 3 TOG molecules (Figure 4C), where the ester moieties potentially coordinated to the metal. Such behavior was described before for phospholipid membranes (45). The RMSD graph shows that there was a change in the system box dimensions for NPT ensemble after ~65 ns (Figure 4D). Analysis of change in TOG core atoms C1-C2-C3 and O11-O21-O31 angles with time suggests that increased variability in the dihedral angles of the TOG backbone underpin this change (Figure 4E). The radius of gyration graph for TOGs also confirmed this observation (Figure 4F). It appears that at ~65 ns the system exits a local energy minimum, leading to substantial re-arrangement. As part of this, DOPC and TOG intercalation increased; by reducing the DOPC density, this structural change is the likely cause of the incursion of water and even hydrated ions towards the core of the 4L system (Supplementary Figure S6).

## Discussion

Computational models has always provided a good starting point to test or conduct an experiment which otherwise was not possible (46). At many places the results obtained from computational models were proved experimentally when experimental techniques were available. The best example is the quantum calculation of Cu-Cu bonds which were later proved by experimentally (47). In principal, if computational model is built accurately, it may provide useful results. MOG forcefield used in current study was calculated through restrained electrostatic potential (RESP) method using Gaussian tool (31, 48, 49). The platform of RESP and ESP charge derive (R.E.D) server was used to build input files and perform QM calculations. There are many methods to calculate forcefields but RESP method implemented by R.E.D server is considered best among which provide charge reproducibility as high as 0.0001 e (33). Due to high compatibility with AMBER, almost all AMBER forcefields utilize RESP method with Gaussian (27). As simulations in current study were intended to run using AMBER, the RESP method using Gaussian were considered most appropriate choice. There were many limitations for this study; primary limitation was the computational power and available tools to prepare membrane systems. Manual membrane building was performed for some experiments due to unavailability of tools offering required functionality. Another limitation was related to TOG molecule building. As when hydrogens were removed from MOG and OL tails were attached to it, the TOG molecule as whole got very small charge of (−0.14). We implemented modular approach of AMBER for building TOG molecule that has enabled us to perform simulations at atomic scale, it is still suggested that some forcefield optimizations in terms of charge are required for TOG molecule. The results of atomic level simulations performed in this study showed many comparable observations to CG simulations.

TAGs are highly dynamic molecules and in some cases have long chains of carbon atoms. These long chains of heavy atoms increase collision and movement rate of atoms causing TAGs to have different conformations at different timescales. One study performed conformational analysis of TAGs through UA simulations (23). During course of simulations it was shown that TAGs acquired different conformations from low angle to high angle variable extended conformations. It is considered that these extended conformations may have functional and metabolic importance. At low temperature TAGs can arrange themselves in layered structures but at physiological temperature simulations it was observed that TAGs acquire many different conformations casting away layered structures. Spontaneous and random aggregation inside ER bilayer also ruled out multilamellar structures of TAGs (14, 20). As properly built system is of the primary importance for a successful simulation, still it is unclear that which TAG conformation would be better to build initial membrane system. The CG and UA simulations has not essentially provided details of initial TAG conformation and even not the number of TAGs according to given conformation. Considering group of atoms as single entity or bead is no doubt a big achievement and is serving as a tool to simulate larger systems (50–52). As a tradeoff, beads do not essentially represent actual volume and molecule to molecule distances. Due to which in most cases number of TOGs are taken based on the system size to fill the space. To deal with these issues atomic level simulations are better because they provide a more acceptable representation of atoms and their bonds. Even when it comes to analogy, atomic level simulations are considered better option due to their contiguousness with real experiments (53, 54). In this study, firstly, we performed atomic level simulation of two most stable conformations of TOGs i.e., TOG3 and TOG2:1 at 273K, 277K, and 298K. During simulation run TOG acquired many conformations but our results demonstrated that TOG2:1 was the most populated conformation at different temperatures and is more favorable choice to build initial membrane system. Secondly, we built TOG-only membranes and simulated varied number of TOGs based on distances between them (12 Å, 6 Å, and 3 Å) in membranes of same area. It was observed that the system with TOG-TOG distance of 6Å had relatively lower and stable RMSD. It was also supported by moderate change in volume and almost unchanged radius of gyration from start to end of simulation.

Many studies provided evidence that LDs may be synthesized inside ER membranes and after reaching certain size LDs can pinch themselves off from ER with the help of proteins most important of which are caveolins. Although the complete mechanism of LD formation and pinching is still to be uncovered but when TAGs were put inside ER mimicking bilayer during CG simulations, TAGs caused positive curvature which was enhanced by addition of truncated cav1 protein (20). It was suggested that TAGs have the ability to aggregate and make TAG lens which with the help of proteins enable LDs to pinch-off from ER. Similarly when put in NVT ensemble, rectangular packed lipids from edges moved upside which caused a curvature leading to LD vesiculation (15). We observed similar behavior when TOGs were simulated with NVT ensemble. TOGs tend to make a curved structure by moving up from edges reaching the volume limit after which the curved structure was deformed. It is currently not understood why TOGs behaved like this under NVT ensemble and may need more experimentations.

A key observation from MD simulation of 4L system suggests that a fraction of Na^+^ ions and water molecules diffused towards core of the 4L membrane (Supplementary Figure S6). It is currently not possible to deduce whether movement/presence of positive ions and water in TOG core is an artefact because there is one such example where through freeze-fracture immunocytochemistry and electron microscopy, authors showed presence of PAT family proteins in LD core (55). Except this in CG simulations water molecule was demonstrated to move from surface to TAG core. Some TAGs were also shown to protrude out to surface reaching up phospholipid monolayer barrier. In another study 6 Na^+^ ions were put in trilayer system (TAG layer at core with phospholipid monolayers) but whether Na^+^ ions interacted with TAGs or not was not discussed (13). But movement of water or ions inside TAG core still needs some explanation. One possibility could be the presence of small charge on TOG molecule which may have created a charge imbalance in the system. Other explanation could be the intercalation of TOGs with DOPC as demonstrated in our and earlier studies. In our study it was observed that TOGs and DOPCs have tail-tail intercalations which caused some deformations in the membrane which may have given space to movement of water molecules with ions inside TOG core. Another argument could be that as compared to bilayer systems, monolayer systems are not that stable and show least level of permeability and less stability. We did not used proteins in current study, it is possible that in nature these proteins may act as permeability agents to control movement of ions and solute across phospholipid membranes. Still, these observations require further experimentation and validation and in future studies we will try to address them.

### TAG Packing Model

Seeds of oil producing plants are rich in oil bodies/LDs while they are also rich in many essential positive ions and other important molecules (56–59). Except neutral lipids LDs also contain many other important nutrients but whether they also contains positive ions is not still clear (3). Even when it comes to use A/LDs as an efficient drug carrier system, it needs to understand how drug molecules can be packed inside A/LD core. For that purpose and based on our understandings of A/LDs, a TAG-packing model was developed.

Under physiological temperature, TAGs are isotropic, phospholipids are anisotropic, cholesterols are liquid crystals, and this all makes overall lipid droplets as semi-isotropic structures (30, 60, 61). Due to isotropic features, TAGs can be considered as homogeneous entity and general gas laws through van der Waals equation of state can be applied on it (62, 63). Considering phospholipids as container where TAGs are needed to be placed, it can be imagined that inner walls of phospholipids are in fact non reacting species i.e. long carbon chains. The neutral nature and isotropic property of TAGs would help TAGs to align themselves inside phospholipid container appropriately. During aligning of TAGs, it can be expected that TAGs would follow gas equations. In this scenario, phospholipids would exert pressure on TAGs and TAGs themselves would also exert pressure on each other. Phospholipids would maintain volume of container and temperature helps in movement of molecules featuring a certain dynamic equilibrium. As it was observed through simulation results that one Na^+^ ion was buried by three TOGs, it would not only effect pressure but also effect volume of system so that possibly groups of TAG molecules that had aligned themselves with positive ions, would act as a unit. These units would definitely make a charge density around them so that one charge density would push away other charge density. Such disturbances would cause discrepancies in packing of TAGs under phospholipid container. It would increase volume which would eventually decrease pressure. According to gas equation, to prevent pressure decrease, number of molecules can be reduced and/or temperature can be reduced. But in case of LDs, cells don’t reduce number of atoms nor temperature; in fact they would rather increase number of TAGs to increase pressure. Hence, to balance the equation, where pressure, temperature, and number of atoms must increase, a stabilizer is required which would at one hand manage charge, at other hand would reduce density. The stabilizer is an entity which will control TAG units eventually causing repulsive interactions to reduce and hydrophobic interactions to take over. The stabilizer could be small molecule like cholesterol, an ion like Na^+^, or a peptide that may maintain neutral charge density.

Phospholipid container is dynamic in terms of the elasticity of phospholipid bonds with each other. It means the volume of phospholipid container may change, but this volume change as compared with volume of molecule it is containing, would be very small. It would make phospholipid container volume more or less constant as change of volume would not be enough to allow a big molecule (like TAG) to enter in the system if system is already full of molecules. Therefore, in case of phospholipid container, in one unit time where if volume remains constant, the pressure needs to be stabilized. In this case volume term in van der Waals equation would not need attention, but the pressure needs to be considered. Van der Waals added (a/V^2^) term to calculate average attractions of particles per volume, while in case of TAGs, attraction of particles would halt up to a limit of charges where TAGs would start pushing each other away. It suggests that for TAG packing, inside phospholipid monolayer, van der Waals equation would require a stabilizer which not only stabilizes charge but also stabilizes the overall density and maintains the constant volume. The stabilizer term for van der Waals pressure could be V/q (charge density) as follows

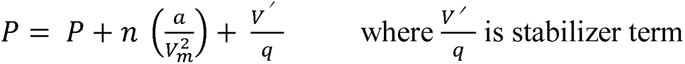

It would modify van der Waals equation as

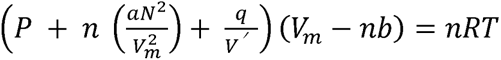

The stabilizer charge *q* would be equal and opposite to charge of attractive forces *a, n* is number of entities in one unit that need stabilizer, and volume of system would decrease to a factor of stabilizer volume *V*’.

