## Supplementary figures and images for "Atomistic Molecular Dynamics Simulations of Trioleoylglycerol – Phospholipid Membrane Systems"

### Supplementary Figure S1

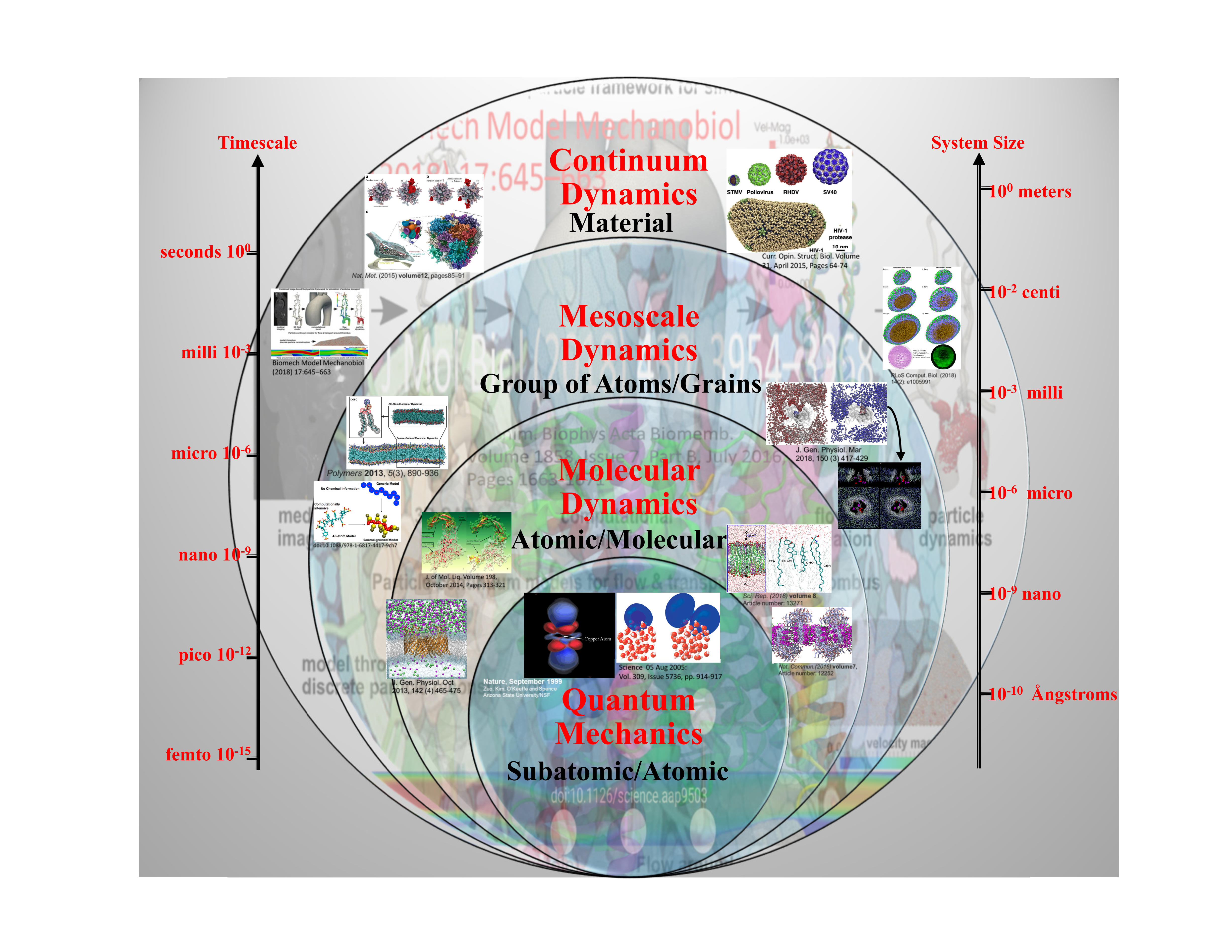

### Supplementary Figure S6

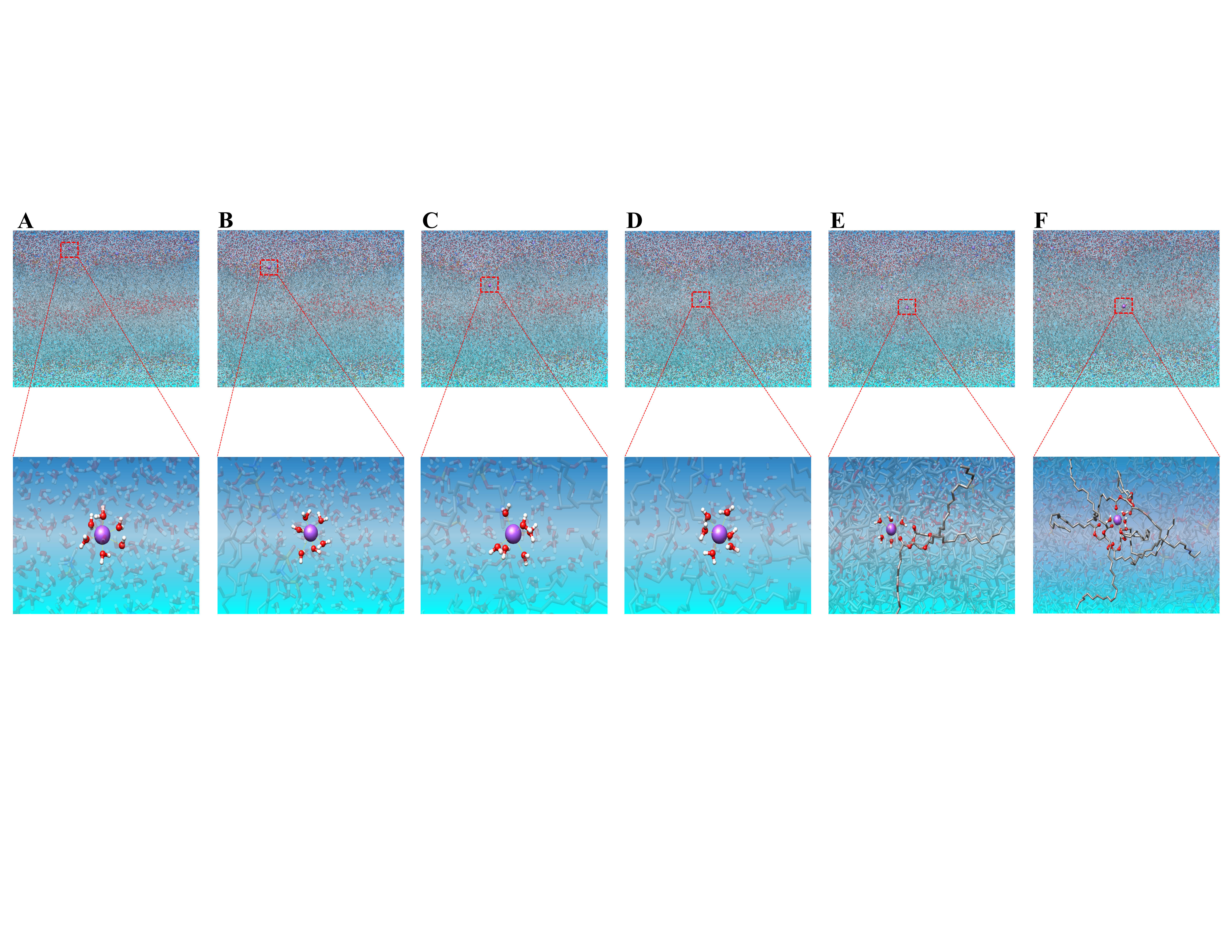
