## Supplementary File S2 for "Atomistic Molecular Dynamics Simulations of Trioleoylglycerol – Phospholipid Membrane Systems"

```
MOG Forcefield

!!index array str
 "MOG"
!entry.MOG.unit.atoms table  str name  str type  int typex  int resx  int flags  int seq  int elmnt  dbl chg
 "C11" "cC" 0 1 131075 1 6 0.474000
 "O12" "oC" 0 1 131075 2 8 -0.474200
 "O11" "oS" 0 1 131075 3 8 -0.235600
 "C1" "cA" 0 1 131075 4 6 -0.023500
 "HR" "hE" 0 1 131075 5 1 0.093000
 "HS" "hE" 0 1 131075 6 1 0.114400
 "C2" "cA" 0 1 131075 7 6 -0.028800
 "HX" "hE" 0 1 131075 8 1 0.171100
 "O21" "oS" 0 1 131075 9 8 -0.272100
 "C21" "cC" 0 1 131075 10 6 0.480500
 "O22" "oC" 0 1 131075 11 8 -0.500000
 "C3" "cA" 0 1 131075 12 6 -0.054300
 "HA" "hE" 0 1 131075 13 1 0.117400
 "HB" "hE" 0 1 131075 14 1 0.129700
 "C12" "cD" 0 1 131075 15 6 0.107600
 "H2R" "hL" 0 1 131075 16 1 0.009700
 "H2S" "hL" 0 1 131075 17 1 -0.021100
 "C13" "cD" 0 1 131075 18 6 -0.028300
 "H3R" "hL" 0 1 131075 19 1 -0.005200
 "H3S" "hL" 0 1 131075 20 1 0.047300
 "C14" "cD" 0 1 131075 21 6 -0.056300
 "H4R" "hL" 0 1 131075 22 1 0.008300
 "H4S" "hL" 0 1 131075 23 1 0.045100
 "C15" "cD" 0 1 131075 24 6 -0.007400
 "H5R" "hL" 0 1 131075 25 1 0.002900
 "H5S" "hL" 0 1 131075 26 1 0.017300
 "C16" "cD" 0 1 131075 27 6 -0.011300
 "H6R" "hL" 0 1 131075 28 1 0.0
 "H6S" "hL" 0 1 131075 29 1 0.0
 "C17" "cD" 0 1 131075 30 6 0.048600
 "H7R" "hL" 0 1 131075 31 1 0.002500
 "H7S" "hL" 0 1 131075 32 1 -0.003700
 "C18" "cD" 0 1 131075 33 6 0.021300
 "H8R" "hL" 0 1 131075 34 1 0.026200
 "H8S" "hL" 0 1 131075 35 1 0.026200
 "C19" "cB" 0 1 131075 36 6 -0.219100
 "H9R" "hB" 0 1 131075 37 1 0.109300
 "C110" "cB" 0 1 131075 38 6 -0.215100
 "H10R" "hB" 0 1 131075 39 1 0.106300
 "C111" "cD" 0 1 131075 40 6 0.047800
 "H11R" "hL" 0 1 131075 41 1 0.026300
 "H11S" "hL" 0 1 131075 42 1 0.026300
 "C112" "cD" 0 1 131075 43 6 -0.001800
 "H12R" "hL" 0 1 131075 44 1 0.007000
 "H12S" "hL" 0 1 131075 45 1 -0.022000
 "C113" "cD" 0 1 131075 46 6 0.064500
 "H13R" "hL" 0 1 131075 47 1 -0.007200
 "H13S" "hL" 0 1 131075 48 1 -0.017500
 "C114" "cD" 0 1 131075 49 6 0.015500
 "H14R" "hL" 0 1 131075 50 1 -0.002800
 "H14S" "hL" 0 1 131075 51 1 -0.002800
 "C115" "cD" 0 1 131075 52 6 0.005500
 "H15R" "hL" 0 1 131075 53 1 -0.011900
 "H15S" "hL" 0 1 131075 54 1 -0.005800
 "C116" "cD" 0 1 131075 55 6 -0.030500
 "H16R" "hL" 0 1 131075 56 1 0.002900
 "H16S" "hL" 0 1 131075 57 1 0.002900
 "C117" "cD" 0 1 131075 58 6 0.113400
 "H17R" "hL" 0 1 131075 59 1 -0.032100
 "H17S" "hL" 0 1 131075 60 1 -0.032100
 "C118" "cD" 0 1 131075 61 6 -0.058700
 "H18R" "hL" 0 1 131075 62 1 0.003900
 "H18S" "hL" 0 1 131075 63 1 0.014700
 "H18T" "hL" 0 1 131075 64 1 0.008700
 "O31" "oS" 0 1 131075 65 8 -0.250900
 "C31" "cC" 0 1 131075 66 6 0.468200
 "O32" "oC" 0 1 131075 67 8 -0.471000
!entry.MOG.unit.atomspertinfo table  str pname  str ptype  int ptypex  int pelmnt  dbl pchg
 "C11" "cC" 0 -1 0.0
 "O12" "oC" 0 -1 0.0
 "O11" "oS" 0 -1 0.0
 "C1" "cA" 0 -1 0.0
 "HR" "hE" 0 -1 0.0
 "HS" "hE" 0 -1 0.0
 "C2" "cA" 0 -1 0.0
 "HX" "hE" 0 -1 0.0
 "O21" "oS" 0 -1 0.0
 "C21" "cC" 0 -1 0.0
 "O22" "oC" 0 -1 0.0
 "C3" "cA" 0 -1 0.0
 "HA" "hE" 0 -1 0.0
 "HB" "hE" 0 -1 0.0
 "C12" "cD" 0 -1 0.0
 "H2R" "hL" 0 -1 0.0
 "H2S" "hL" 0 -1 0.0
 "C13" "cD" 0 -1 0.0
 "H3R" "hL" 0 -1 0.0
 "H3S" "hL" 0 -1 0.0
 "C14" "cD" 0 -1 0.0
 "H4R" "hL" 0 -1 0.0
 "H4S" "hL" 0 -1 0.0
 "C15" "cD" 0 -1 0.0
 "H5R" "hL" 0 -1 0.0
 "H5S" "hL" 0 -1 0.0
 "C16" "cD" 0 -1 0.0
 "H6R" "hL" 0 -1 0.0
 "H6S" "hL" 0 -1 0.0
 "C17" "cD" 0 -1 0.0
 "H7R" "hL" 0 -1 0.0
 "H7S" "hL" 0 -1 0.0
 "C18" "cD" 0 -1 0.0
 "H8R" "hL" 0 -1 0.0
 "H8S" "hL" 0 -1 0.0
 "C19" "cB" 0 -1 0.0
 "H9R" "hB" 0 -1 0.0
 "C110" "cB" 0 -1 0.0
 "H10R" "hB" 0 -1 0.0
 "C111" "cD" 0 -1 0.0
 "H11R" "hL" 0 -1 0.0
 "H11S" "hL" 0 -1 0.0
 "C112" "cD" 0 -1 0.0
 "H12R" "hL" 0 -1 0.0
 "H12S" "hL" 0 -1 0.0
 "C113" "cD" 0 -1 0.0
 "H13R" "hL" 0 -1 0.0
 "H13S" "hL" 0 -1 0.0
 "C114" "cD" 0 -1 0.0
 "H14R" "hL" 0 -1 0.0
 "H14S" "hL" 0 -1 0.0
 "C115" "cD" 0 -1 0.0
 "H15R" "hL" 0 -1 0.0
 "H15S" "hL" 0 -1 0.0
 "C116" "cD" 0 -1 0.0
 "H16R" "hL" 0 -1 0.0
 "H16S" "hL" 0 -1 0.0
 "C117" "cD" 0 -1 0.0
 "H17R" "hL" 0 -1 0.0
 "H17S" "hL" 0 -1 0.0
 "C118" "cD" 0 -1 0.0
 "H18R" "hL" 0 -1 0.0
 "H18S" "hL" 0 -1 0.0
 "H18T" "hL" 0 -1 0.0
 "O31" "oS" 0 -1 0.0
 "C31" "cC" 0 -1 0.0
 "O32" "oC" 0 -1 0.0
!entry.MOG.unit.boundbox array dbl
 -1.000000
 0.0
 0.0
 0.0
 0.0
!entry.MOG.unit.childsequence single int
 2
!entry.MOG.unit.connect array int
 1
 66
!entry.MOG.unit.connectivity table  int atom1x  int atom2x  int flags
 1 2 2
 1 3 1
 3 4 1
 4 5 1
 4 6 1
 4 7 1
 7 8 1
 7 9 1
 7 12 1
 9 10 1
 10 11 2
 10 15 1
 12 13 1
 12 14 1
 12 65 1
 15 16 1
 15 17 1
 15 18 1
 18 19 1
 18 20 1
 18 21 1
 21 22 1
 21 23 1
 21 24 1
 24 25 1
 24 26 1
 24 27 1
 27 28 1
 27 29 1
 27 30 1
 30 31 1
 30 32 1
 30 33 1
 33 34 1
 33 35 1
 33 36 1
 36 37 1
 36 38 2
 38 39 1
 38 40 1
 40 41 1
 40 42 1
 40 43 1
 43 44 1
 43 45 1
 43 46 1
 46 47 1
 46 48 1
 46 49 1
 49 50 1
 49 51 1
 49 52 1
 52 53 1
 52 54 1
 52 55 1
 55 56 1
 55 57 1
 55 58 1
 58 59 1
 58 60 1
 58 61 1
 61 62 1
 61 63 1
 61 64 1
 65 66 1
 66 67 2
!entry.MOG.unit.hierarchy table  str abovetype  int abovex  str belowtype  int belowx
 "U" 0 "R" 1
 "R" 1 "A" 1
 "R" 1 "A" 2
 "R" 1 "A" 3
 "R" 1 "A" 4
 "R" 1 "A" 5
 "R" 1 "A" 6
 "R" 1 "A" 7
 "R" 1 "A" 8
 "R" 1 "A" 9
 "R" 1 "A" 10
 "R" 1 "A" 11
 "R" 1 "A" 12
 "R" 1 "A" 13
 "R" 1 "A" 14
 "R" 1 "A" 15
 "R" 1 "A" 16
 "R" 1 "A" 17
 "R" 1 "A" 18
 "R" 1 "A" 19
 "R" 1 "A" 20
 "R" 1 "A" 21
 "R" 1 "A" 22
 "R" 1 "A" 23
 "R" 1 "A" 24
 "R" 1 "A" 25
 "R" 1 "A" 26
 "R" 1 "A" 27
 "R" 1 "A" 28
 "R" 1 "A" 29
 "R" 1 "A" 30
 "R" 1 "A" 31
 "R" 1 "A" 32
 "R" 1 "A" 33
 "R" 1 "A" 34
 "R" 1 "A" 35
 "R" 1 "A" 36
 "R" 1 "A" 37
 "R" 1 "A" 38
 "R" 1 "A" 39
 "R" 1 "A" 40
 "R" 1 "A" 41
 "R" 1 "A" 42
 "R" 1 "A" 43
 "R" 1 "A" 44
 "R" 1 "A" 45
 "R" 1 "A" 46
 "R" 1 "A" 47
 "R" 1 "A" 48
 "R" 1 "A" 49
 "R" 1 "A" 50
 "R" 1 "A" 51
 "R" 1 "A" 52
 "R" 1 "A" 53
 "R" 1 "A" 54
 "R" 1 "A" 55
 "R" 1 "A" 56
 "R" 1 "A" 57
 "R" 1 "A" 58
 "R" 1 "A" 59
 "R" 1 "A" 60
 "R" 1 "A" 61
 "R" 1 "A" 62
 "R" 1 "A" 63
 "R" 1 "A" 64
 "R" 1 "A" 65
 "R" 1 "A" 66
 "R" 1 "A" 67
!entry.MOG.unit.name single str
 "MOG"
!entry.MOG.unit.positions table  dbl x  dbl y  dbl z
 -6.433977 -3.294948 -0.465815
 -5.916197 -4.235186 0.010898
 -5.742159 -2.337027 -1.083473
 -6.409521 -1.254334 -1.686022
 -7.479119 -1.316924 -1.514082
 -6.223297 -1.297121 -2.752628
 -5.871467 0.041211 -1.101757
 -4.800774 0.083976 -1.239125
 -6.224968 0.111112 0.258398
 -5.453488 -0.266554 1.307345
 -5.990993 -0.396105 2.351171
 -6.497678 1.253594 -1.770585
 -7.568638 1.263114 -1.593163
 -6.324730 1.229816 -2.838310
 -3.962297 -0.425593 1.118974
 -3.634324 -1.002141 1.971551
 -3.737730 -1.002743 0.230491
 -3.249731 0.939279 1.086220
 -3.664707 1.556918 0.295479
 -3.455192 1.460311 2.017394
 -1.735249 0.830257 0.878037
 -1.542285 0.321488 -0.063882
 -1.342297 1.838399 0.765735
 -0.986462 0.132651 2.018397
 -1.295817 0.580111 2.961135
 -1.273346 -0.915448 2.069905
 0.540607 0.222345 1.917181
 0.837367 1.269906 1.885999
 0.967815 -0.186092 2.830917
 1.148172 -0.513450 0.720411
 0.811550 -1.549444 0.726032
 0.793141 -0.080870 -0.211071
 2.682621 -0.479758 0.721926
 3.020543 0.550493 0.713064
 3.042021 -0.912590 1.654841
 3.273865 -1.255398 -0.425950
 3.053727 -2.313107 -0.409206
 4.002131 -0.808478 -1.435719
 4.328683 -1.536390 -2.164925
 4.450592 0.600405 -1.727764
 4.030917 0.896462 -2.688209
 4.057999 1.297557 -0.995618
 5.978464 0.746372 -1.804636
 6.368873 0.039626 -2.535682
 6.212384 1.738328 -2.186221
 6.691254 0.542223 -0.467012
 6.300246 1.255828 0.257569
 6.457163 -0.445619 -0.077552
 8.209530 0.703522 -0.565497
 8.591833 -0.000988 -1.301057
 8.443710 1.697566 -0.945648
 8.918532 0.496295 0.775178
 8.472299 1.162169 1.511665
 8.724751 -0.512708 1.132467
 10.428863 0.764052 0.751201
 10.593805 1.800944 0.463070
 10.810779 0.669153 1.766425
 11.264657 -0.134977 -0.168983
 10.958301 0.001958 -1.202939
 12.297860 0.201336 -0.118250
 11.214474 -1.622251 0.183084
 11.891205 -2.190816 -0.447649
 10.220557 -2.037636 0.050491
 11.507067 -1.789817 1.216434
 -5.900566 2.430387 -1.291329
 -6.471662 3.090150 -0.281983
 -5.962102 4.021800 0.222291
!entry.MOG.unit.residueconnect table  int c1x  int c2x  int c3x  int c4x  int c5x  int c6x
 0 0 0 0 0 0
!entry.MOG.unit.residues table  str name  int seq  int childseq  int startatomx  str restype  int imagingx
 "MOG" 1 70 1 "?" 0
!entry.MOG.unit.residuesPdbSequenceNumber array int
 0
!entry.MOG.unit.solventcap array dbl
 -1.000000
 0.0
 0.0
 0.0
 0.0
!entry.MOG.unit.velocities table  dbl x  dbl y  dbl z
 0.0 0.0 0.0
 0.0 0.0 0.0
 0.0 0.0 0.0
 0.0 0.0 0.0
 0.0 0.0 0.0
 0.0 0.0 0.0
 0.0 0.0 0.0
 0.0 0.0 0.0
 0.0 0.0 0.0
 0.0 0.0 0.0
 0.0 0.0 0.0
 0.0 0.0 0.0
 0.0 0.0 0.0
 0.0 0.0 0.0
 0.0 0.0 0.0
 0.0 0.0 0.0
 0.0 0.0 0.0
 0.0 0.0 0.0
 0.0 0.0 0.0
 0.0 0.0 0.0
 0.0 0.0 0.0
 0.0 0.0 0.0
 0.0 0.0 0.0
 0.0 0.0 0.0
 0.0 0.0 0.0
 0.0 0.0 0.0
 0.0 0.0 0.0
 0.0 0.0 0.0
 0.0 0.0 0.0
 0.0 0.0 0.0
 0.0 0.0 0.0
 0.0 0.0 0.0
 0.0 0.0 0.0
 0.0 0.0 0.0
 0.0 0.0 0.0
 0.0 0.0 0.0
 0.0 0.0 0.0
 0.0 0.0 0.0
 0.0 0.0 0.0
 0.0 0.0 0.0
 0.0 0.0 0.0
 0.0 0.0 0.0
 0.0 0.0 0.0
 0.0 0.0 0.0
 0.0 0.0 0.0
 0.0 0.0 0.0
 0.0 0.0 0.0
 0.0 0.0 0.0
 0.0 0.0 0.0
 0.0 0.0 0.0
 0.0 0.0 0.0
 0.0 0.0 0.0
 0.0 0.0 0.0
 0.0 0.0 0.0
 0.0 0.0 0.0
 0.0 0.0 0.0
 0.0 0.0 0.0
 0.0 0.0 0.0
 0.0 0.0 0.0
 0.0 0.0 0.0
 0.0 0.0 0.0
 0.0 0.0 0.0
 0.0 0.0 0.0
 0.0 0.0 0.0
 0.0 0.0 0.0
 0.0 0.0 0.0
 0.0 0.0 0.0

TOG2:1 PDB File

HETATM    1  C12 OL  A   1      33.015  53.926  27.596  1.00  0.00           C
HETATM    2  H2R OL  A   1      32.915  52.847  27.463  1.00  0.00           H
HETATM    3  H2S OL  A   1      33.840  54.201  26.935  1.00  0.00           H
HETATM    4  C13 OL  A   1      33.368  54.394  29.019  1.00  0.00           C
HETATM    5  H3R OL  A   1      33.411  55.483  29.088  1.00  0.00           H
HETATM    6  H3S OL  A   1      34.365  54.027  29.271  1.00  0.00           H
HETATM    7  C14 OL  A   1      32.462  53.871  30.148  1.00  0.00           C
HETATM    8  H4R OL  A   1      31.471  54.321  30.243  1.00  0.00           H
HETATM    9  H4S OL  A   1      32.180  52.827  29.997  1.00  0.00           H
HETATM   10  C15 OL  A   1      32.992  54.076  31.559  1.00  0.00           C
HETATM   11  H5R OL  A   1      33.243  55.121  31.755  1.00  0.00           H
HETATM   12  H5S OL  A   1      33.956  53.570  31.648  1.00  0.00           H
HETATM   13  C16 OL  A   1      32.121  53.603  32.732  1.00  0.00           C
HETATM   14  H6R OL  A   1      31.289  54.310  32.758  1.00  0.00           H
HETATM   15  H6S OL  A   1      31.690  52.627  32.499  1.00  0.00           H
HETATM   16  C17 OL  A   1      32.819  53.658  34.110  1.00  0.00           C
HETATM   17  H7R OL  A   1      33.054  54.711  34.278  1.00  0.00           H
HETATM   18  H7S OL  A   1      33.759  53.102  34.143  1.00  0.00           H
HETATM   19  C18 OL  A   1      31.935  53.092  35.189  1.00  0.00           C
HETATM   20  H8R OL  A   1      30.999  53.653  35.151  1.00  0.00           H
HETATM   21  H8S OL  A   1      31.615  52.078  34.940  1.00  0.00           H
HETATM   22  C19 OL  A   1      32.541  53.163  36.551  1.00  0.00           C
HETATM   23  H9R OL  A   1      33.591  53.442  36.591  1.00  0.00           H
HETATM   24 C110 OL  A   1      31.988  52.754  37.684  1.00  0.00           C
HETATM   25 H10R OL  A   1      30.942  52.458  37.676  1.00  0.00           H
HETATM   26 C111 OL  A   1      32.751  52.566  38.974  1.00  0.00           C
HETATM   27 H11R OL  A   1      33.496  53.350  39.124  1.00  0.00           H
HETATM   28 H11S OL  A   1      33.142  51.548  39.017  1.00  0.00           H
HETATM   29 C112 OL  A   1      31.806  52.720  40.111  1.00  0.00           C
HETATM   30 H12R OL  A   1      31.306  53.690  40.092  1.00  0.00           H
HETATM   31 H12S OL  A   1      30.937  52.072  39.977  1.00  0.00           H
HETATM   32 C113 OL  A   1      32.443  52.391  41.488  1.00  0.00           C
HETATM   33 H13R OL  A   1      33.452  52.796  41.592  1.00  0.00           H
HETATM   34 H13S OL  A   1      32.616  51.333  41.692  1.00  0.00           H
HETATM   35 C114 OL  A   1      31.628  52.846  42.653  1.00  0.00           C
HETATM   36 H14R OL  A   1      31.422  53.918  42.648  1.00  0.00           H
HETATM   37 H14S OL  A   1      30.688  52.290  42.644  1.00  0.00           H
HETATM   38 C115 OL  A   1      32.251  52.515  44.005  1.00  0.00           C
HETATM   39 H15R OL  A   1      32.999  53.275  44.237  1.00  0.00           H
HETATM   40 H15S OL  A   1      32.617  51.487  44.048  1.00  0.00           H
HETATM   41 C116 OL  A   1      31.186  52.616  45.106  1.00  0.00           C
HETATM   42 H16R OL  A   1      30.741  53.592  44.898  1.00  0.00           H
HETATM   43 H16S OL  A   1      30.422  51.838  45.052  1.00  0.00           H
HETATM   44 C117 OL  A   1      31.789  52.536  46.441  1.00  0.00           C
HETATM   45 H17R OL  A   1      32.654  53.183  46.599  1.00  0.00           H
HETATM   46 H17S OL  A   1      32.121  51.522  46.670  1.00  0.00           H
HETATM   47 C118 OL  A   1      30.747  52.848  47.521  1.00  0.00           C
HETATM   48 H18R OL  A   1      30.228  53.790  47.327  1.00  0.00           H
HETATM   49 H18S OL  A   1      29.979  52.072  47.551  1.00  0.00           H
HETATM   50 H18T OL  A   1      31.245  52.816  48.492  1.00  0.00           H
TER
HETATM   51  C11 MOG A   1      31.744  54.579  27.078  1.00  0.00           C
HETATM   52  O12 MOG A   1      30.890  55.111  27.736  1.00  0.00           O
HETATM   53  O11 MOG A   1      31.686  54.367  25.763  1.00  0.00           O
HETATM   54  C1  MOG A   1      30.475  54.745  25.019  1.00  0.00           C
HETATM   55  HR  MOG A   1      29.540  54.488  25.522  1.00  0.00           H
HETATM   56  HS  MOG A   1      30.485  55.838  25.016  1.00  0.00           H
HETATM   57  C2  MOG A   1      30.469  54.278  23.537  1.00  0.00           C
HETATM   58  HX  MOG A   1      31.454  54.540  23.143  1.00  0.00           H
HETATM   59  O21 MOG A   1      29.435  54.911  22.720  1.00  0.00           O
HETATM   60  C21 MOG A   1      29.586  54.929  21.349  1.00  0.00           C
HETATM   61  O22 MOG A   1      30.605  54.660  20.760  1.00  0.00           O
HETATM   62  C3  MOG A   1      30.291  52.731  23.401  1.00  0.00           C
HETATM   63  HA  MOG A   1      29.258  52.546  23.707  1.00  0.00           H
HETATM   64  HB  MOG A   1      30.409  52.320  22.395  1.00  0.00           H
HETATM   65  C12 MOG A   1      28.332  55.581  20.813  1.00  0.00           C
HETATM   66  H2R MOG A   1      27.430  55.199  21.297  1.00  0.00           H
HETATM   67  H2S MOG A   1      28.266  56.643  21.058  1.00  0.00           H
HETATM   68  C13 MOG A   1      28.168  55.397  19.312  1.00  0.00           C
HETATM   69  H3R MOG A   1      28.014  54.338  19.093  1.00  0.00           H
HETATM   70  H3S MOG A   1      29.074  55.652  18.760  1.00  0.00           H
HETATM   71  C14 MOG A   1      26.962  56.265  18.785  1.00  0.00           C
HETATM   72  H4R MOG A   1      26.029  55.872  19.193  1.00  0.00           H
HETATM   73  H4S MOG A   1      27.054  57.289  19.156  1.00  0.00           H
HETATM   74  C15 MOG A   1      26.759  56.383  17.237  1.00  0.00           C
HETATM   75  H5R MOG A   1      25.788  56.850  17.062  1.00  0.00           H
HETATM   76  H5S MOG A   1      26.602  55.370  16.860  1.00  0.00           H
HETATM   77  C16 MOG A   1      27.832  57.068  16.400  1.00  0.00           C
HETATM   78  H6R MOG A   1      27.948  58.084  16.784  1.00  0.00           H
HETATM   79  H6S MOG A   1      28.810  56.670  16.679  1.00  0.00           H
HETATM   80  C17 MOG A   1      27.618  56.932  14.897  1.00  0.00           C
HETATM   81  H7R MOG A   1      26.634  57.319  14.627  1.00  0.00           H
HETATM   82  H7S MOG A   1      27.546  55.850  14.769  1.00  0.00           H
HETATM   83  C18 MOG A   1      28.749  57.582  14.102  1.00  0.00           C
HETATM   84  H8R MOG A   1      28.660  58.657  14.275  1.00  0.00           H
HETATM   85  H8S MOG A   1      29.730  57.253  14.451  1.00  0.00           H
HETATM   86  C19 MOG A   1      28.601  57.203  12.679  1.00  0.00           C
HETATM   87  H9R MOG A   1      27.605  57.278  12.249  1.00  0.00           H
HETATM   88 C110 MOG A   1      29.587  56.781  11.940  1.00  0.00           C
HETATM   89 H10R MOG A   1      30.607  56.702  12.309  1.00  0.00           H
HETATM   90 C111 MOG A   1      29.336  56.463  10.517  1.00  0.00           C
HETATM   91 H11R MOG A   1      28.292  56.571  10.213  1.00  0.00           H
HETATM   92 H11S MOG A   1      29.459  55.379  10.478  1.00  0.00           H
HETATM   93 C112 MOG A   1      30.295  57.078   9.531  1.00  0.00           C
HETATM   94 H12R MOG A   1      29.987  58.122   9.442  1.00  0.00           H
HETATM   95 H12S MOG A   1      31.254  57.146  10.050  1.00  0.00           H
HETATM   96 C113 MOG A   1      30.305  56.419   8.115  1.00  0.00           C
HETATM   97 H13R MOG A   1      29.288  56.467   7.722  1.00  0.00           H
HETATM   98 H13S MOG A   1      30.503  55.348   8.195  1.00  0.00           H
HETATM   99 C114 MOG A   1      31.320  57.035   7.129  1.00  0.00           C
HETATM  100 H14R MOG A   1      31.126  58.081   6.882  1.00  0.00           H
HETATM  101 H14S MOG A   1      32.283  56.919   7.630  1.00  0.00           H
HETATM  102 C115 MOG A   1      31.328  56.235   5.825  1.00  0.00           C
HETATM  103 H15R MOG A   1      30.371  56.272   5.300  1.00  0.00           H
HETATM  104 H15S MOG A   1      31.410  55.174   6.070  1.00  0.00           H
HETATM  105 C116 MOG A   1      32.565  56.659   4.989  1.00  0.00           C
HETATM  106 H16R MOG A   1      32.399  57.713   4.757  1.00  0.00           H
HETATM  107 H16S MOG A   1      33.459  56.706   5.615  1.00  0.00           H
HETATM  108 C117 MOG A   1      32.689  55.934   3.730  1.00  0.00           C
HETATM  109 H17R MOG A   1      32.552  54.867   3.917  1.00  0.00           H
HETATM  110 H17S MOG A   1      33.707  56.139   3.392  1.00  0.00           H
HETATM  111 C118 MOG A   1      31.776  56.393   2.567  1.00  0.00           C
HETATM  112 H18R MOG A   1      30.712  56.326   2.806  1.00  0.00           H
HETATM  113 H18S MOG A   1      32.000  57.415   2.256  1.00  0.00           H
HETATM  114 H18T MOG A   1      31.985  55.805   1.671  1.00  0.00           H
HETATM  115  O31 MOG A   1      31.275  52.065  24.193  1.00  0.00           O
HETATM  116  C31 MOG A   1      31.151  50.781  24.485  1.00  0.00           C
HETATM  117  O32 MOG A   1      30.230  50.056  24.120  1.00  0.00           O
TER
HETATM  118  C12 OL  A   1      32.342  50.207  25.261  1.00  0.00           C
HETATM  119  H2R OL  A   1      33.198  50.184  24.584  1.00  0.00           H
HETATM  120  H2S OL  A   1      32.605  50.895  26.067  1.00  0.00           H
HETATM  121  C13 OL  A   1      32.136  48.807  25.898  1.00  0.00           C
HETATM  122  H3R OL  A   1      33.074  48.344  26.209  1.00  0.00           H
HETATM  123  H3S OL  A   1      31.768  48.149  25.108  1.00  0.00           H
HETATM  124  C14 OL  A   1      31.249  48.758  27.091  1.00  0.00           C
HETATM  125  H4R OL  A   1      31.856  49.300  27.819  1.00  0.00           H
HETATM  126  H4S OL  A   1      30.311  49.303  26.965  1.00  0.00           H
HETATM  127  C15 OL  A   1      30.957  47.373  27.650  1.00  0.00           C
HETATM  128  H5R OL  A   1      31.880  46.795  27.720  1.00  0.00           H
HETATM  129  H5S OL  A   1      30.368  46.928  26.846  1.00  0.00           H
HETATM  130  C16 OL  A   1      30.331  47.373  29.024  1.00  0.00           C
HETATM  131  H6R OL  A   1      31.036  47.836  29.718  1.00  0.00           H
HETATM  132  H6S OL  A   1      29.404  47.945  29.103  1.00  0.00           H
HETATM  133  C17 OL  A   1      29.918  46.030  29.520  1.00  0.00           C
HETATM  134  H7R OL  A   1      30.843  45.450  29.547  1.00  0.00           H
HETATM  135  H7S OL  A   1      29.280  45.617  28.737  1.00  0.00           H
HETATM  136  C18 OL  A   1      29.199  46.002  30.881  1.00  0.00           C
HETATM  137  H8R OL  A   1      28.484  46.825  30.937  1.00  0.00           H
HETATM  138  H8S OL  A   1      28.742  45.010  30.884  1.00  0.00           H
HETATM  139  C19 OL  A   1      30.209  46.251  31.923  1.00  0.00           C
HETATM  140  H9R OL  A   1      30.631  47.250  31.847  1.00  0.00           H
HETATM  141 C110 OL  A   1      30.599  45.393  32.841  1.00  0.00           C
HETATM  142 H10R OL  A   1      30.115  44.420  32.822  1.00  0.00           H
HETATM  143 C111 OL  A   1      31.629  45.610  33.915  1.00  0.00           C
HETATM  144 H11R OL  A   1      32.005  46.635  33.933  1.00  0.00           H
HETATM  145 H11S OL  A   1      32.478  44.926  33.875  1.00  0.00           H
HETATM  146 C112 OL  A   1      31.017  45.360  35.298  1.00  0.00           C
HETATM  147 H12R OL  A   1      30.393  46.223  35.538  1.00  0.00           H
HETATM  148 H12S OL  A   1      30.364  44.494  35.428  1.00  0.00           H
HETATM  149 C113 OL  A   1      32.102  45.217  36.379  1.00  0.00           C
HETATM  150 H13R OL  A   1      32.777  46.071  36.297  1.00  0.00           H
HETATM  151 H13S OL  A   1      32.715  44.338  36.172  1.00  0.00           H
HETATM  152 C114 OL  A   1      31.614  45.023  37.791  1.00  0.00           C
HETATM  153 H14R OL  A   1      31.055  45.921  38.062  1.00  0.00           H
HETATM  154 H14S OL  A   1      30.838  44.260  37.887  1.00  0.00           H
HETATM  155 C115 OL  A   1      32.770  44.925  38.818  1.00  0.00           C
HETATM  156 H15R OL  A   1      33.275  45.892  38.863  1.00  0.00           H
HETATM  157 H15S OL  A   1      33.527  44.273  38.378  1.00  0.00           H
HETATM  158 C116 OL  A   1      32.316  44.438  40.203  1.00  0.00           C
HETATM  159 H16R OL  A   1      31.570  45.149  40.565  1.00  0.00           H
HETATM  160 H16S OL  A   1      31.946  43.418  40.090  1.00  0.00           H
HETATM  161 C117 OL  A   1      33.486  44.526  41.192  1.00  0.00           C
HETATM  162 H17R OL  A   1      33.698  45.588  41.332  1.00  0.00           H
HETATM  163 H17S OL  A   1      34.366  44.058  40.745  1.00  0.00           H
HETATM  164 C118 OL  A   1      33.153  43.897  42.548  1.00  0.00           C
HETATM  165 H18R OL  A   1      32.219  44.271  42.972  1.00  0.00           H
HETATM  166 H18S OL  A   1      32.934  42.836  42.408  1.00  0.00           H
HETATM  167 H18T OL  A   1      33.988  43.902  43.253  1.00  0.00           H
TER
END

TOG3 PDB File

HETATM    1  C12 OL    1        56.037  51.289  16.444  1.00  0.00           C
HETATM    2  H2R OL    1        57.096  51.518  16.337  1.00  0.00           H
HETATM    3  H2S OL    1        55.499  52.243  16.539  1.00  0.00           H
HETATM    4  C13 OL    1        55.842  50.405  17.675  1.00  0.00           C
HETATM    5  H3R OL    1        56.611  49.649  17.866  1.00  0.00           H
HETATM    6  H3S OL    1        54.902  49.859  17.564  1.00  0.00           H
HETATM    7  C14 OL    1        55.926  51.244  18.962  1.00  0.00           C
HETATM    8  H4R OL    1        56.981  51.452  19.157  1.00  0.00           H
HETATM    9  H4S OL    1        55.388  52.199  18.937  1.00  0.00           H
HETATM   10  C15 OL    1        55.338  50.473  20.148  1.00  0.00           C
HETATM   11  H5R OL    1        55.646  49.429  20.073  1.00  0.00           H
HETATM   12  H5S OL    1        54.256  50.474  20.187  1.00  0.00           H
HETATM   13  C16 OL    1        55.869  50.990  21.504  1.00  0.00           C
HETATM   14  H6R OL    1        56.962  51.033  21.522  1.00  0.00           H
HETATM   15  H6S OL    1        55.485  51.978  21.767  1.00  0.00           H
HETATM   16  C17 OL    1        55.417  50.096  22.674  1.00  0.00           C
HETATM   17  H7R OL    1        55.673  49.054  22.473  1.00  0.00           H
HETATM   18  H7S OL    1        54.360  50.161  22.941  1.00  0.00           H
HETATM   19  C18 OL    1        56.323  50.343  23.883  1.00  0.00           C
HETATM   20  H8R OL    1        57.195  49.699  23.922  1.00  0.00           H
HETATM   21  H8S OL    1        56.601  51.401  23.873  1.00  0.00           H
HETATM   22  C19 OL    1        55.558  50.022  25.155  1.00  0.00           C
HETATM   23  H9R OL    1        55.215  49.002  25.288  1.00  0.00           H
HETATM   24 C110 OL    1        55.396  50.948  26.086  1.00  0.00           C
HETATM   25 H10R OL    1        55.679  51.965  25.837  1.00  0.00           H
HETATM   26 C111 OL    1        54.647  50.885  27.398  1.00  0.00           C
HETATM   27 H11R OL    1        53.788  50.237  27.244  1.00  0.00           H
HETATM   28 H11S OL    1        54.462  51.876  27.817  1.00  0.00           H
HETATM   29 C112 OL    1        55.543  50.266  28.478  1.00  0.00           C
HETATM   30 H12R OL    1        55.341  49.199  28.563  1.00  0.00           H
HETATM   31 H12S OL    1        56.599  50.323  28.218  1.00  0.00           H
HETATM   32 C113 OL    1        55.181  50.747  29.892  1.00  0.00           C
HETATM   33 H13R OL    1        54.294  50.230  30.278  1.00  0.00           H
HETATM   34 H13S OL    1        54.862  51.784  29.928  1.00  0.00           H
HETATM   35 C114 OL    1        56.307  50.575  30.919  1.00  0.00           C
HETATM   36 H14R OL    1        56.606  49.534  31.026  1.00  0.00           H
HETATM   37 H14S OL    1        57.149  51.134  30.505  1.00  0.00           H
HETATM   38 C115 OL    1        55.972  51.150  32.297  1.00  0.00           C
HETATM   39 H15R OL    1        54.973  50.850  32.613  1.00  0.00           H
HETATM   40 H15S OL    1        55.992  52.245  32.316  1.00  0.00           H
HETATM   41 C116 OL    1        56.915  50.529  33.334  1.00  0.00           C
HETATM   42 H16R OL    1        56.669  49.479  33.481  1.00  0.00           H
HETATM   43 H16S OL    1        57.905  50.689  32.904  1.00  0.00           H
HETATM   44 C117 OL    1        56.722  51.230  34.687  1.00  0.00           C
HETATM   45 H17R OL    1        56.495  50.599  35.550  1.00  0.00           H
HETATM   46 H17S OL    1        55.964  52.015  34.704  1.00  0.00           H
HETATM   47 C118 OL    1        58.129  51.799  34.944  1.00  0.00           C
HETATM   48 H18R OL    1        58.771  50.918  34.939  1.00  0.00           H
HETATM   49 H18S OL    1        58.491  52.439  34.141  1.00  0.00           H
HETATM   50 H18T OL    1        58.150  52.253  35.939  1.00  0.00           H
TER
HETATM   51  C11 MOG   1        55.614  50.677  15.123  1.00  0.00           C
HETATM   52  O12 MOG   1        55.912  49.568  14.740  1.00  0.00           O
HETATM   53  O11 MOG   1        54.836  51.593  14.514  1.00  0.00           O
HETATM   54  C1  MOG   1        53.738  51.167  13.678  1.00  0.00           C
HETATM   55  HR  MOG   1        53.774  51.881  12.857  1.00  0.00           H
HETATM   56  HS  MOG   1        54.020  50.221  13.200  1.00  0.00           H
HETATM   57  C2  MOG   1        52.385  51.345  14.374  1.00  0.00           C
HETATM   58  HX  MOG   1        52.149  52.391  14.563  1.00  0.00           H
HETATM   59  O21 MOG   1        52.572  50.709  15.647  1.00  0.00           O
HETATM   60  C21 MOG   1        51.809  51.094  16.693  1.00  0.00           C
HETATM   61  O22 MOG   1        51.109  52.082  16.628  1.00  0.00           O
HETATM   62  C3  MOG   1        51.161  50.704  13.705  1.00  0.00           C
HETATM   63  HA  MOG   1        51.067  50.961  12.639  1.00  0.00           H
HETATM   64  HB  MOG   1        51.373  49.638  13.758  1.00  0.00           H
HETATM   65  C12 MOG   1        51.821  50.078  17.803  1.00  0.00           C
HETATM   66  H2R MOG   1        51.243  49.216  17.456  1.00  0.00           H
HETATM   67  H2S MOG   1        52.839  49.817  18.099  1.00  0.00           H
HETATM   68  C13 MOG   1        51.204  50.637  19.082  1.00  0.00           C
HETATM   69  H3R MOG   1        50.288  51.178  18.849  1.00  0.00           H
HETATM   70  H3S MOG   1        51.916  51.371  19.484  1.00  0.00           H
HETATM   71  C14 MOG   1        50.710  49.629  20.129  1.00  0.00           C
HETATM   72  H4R MOG   1        49.774  49.226  19.731  1.00  0.00           H
HETATM   73  H4S MOG   1        51.528  48.920  20.285  1.00  0.00           H
HETATM   74  C15 MOG   1        50.450  50.430  21.403  1.00  0.00           C
HETATM   75  H5R MOG   1        49.682  51.129  21.063  1.00  0.00           H
HETATM   76  H5S MOG   1        51.390  50.913  21.677  1.00  0.00           H
HETATM   77  C16 MOG   1        50.047  49.498  22.551  1.00  0.00           C
HETATM   78  H6R MOG   1        49.056  49.077  22.339  1.00  0.00           H
HETATM   79  H6S MOG   1        50.792  48.713  22.655  1.00  0.00           H
HETATM   80  C17 MOG   1        49.984  50.367  23.818  1.00  0.00           C
HETATM   81  H7R MOG   1        49.234  51.125  23.585  1.00  0.00           H
HETATM   82  H7S MOG   1        50.935  50.832  24.072  1.00  0.00           H
HETATM   83  C18 MOG   1        49.501  49.500  24.989  1.00  0.00           C
HETATM   84  H8R MOG   1        48.561  48.954  24.888  1.00  0.00           H
HETATM   85  H8S MOG   1        50.291  48.770  25.209  1.00  0.00           H
HETATM   86  C19 MOG   1        49.383  50.479  26.126  1.00  0.00           C
HETATM   87  H9R MOG   1        48.583  51.200  25.988  1.00  0.00           H
HETATM   88 C110 MOG   1        50.183  50.492  27.180  1.00  0.00           C
HETATM   89 H10R MOG   1        50.985  49.772  27.348  1.00  0.00           H
HETATM   90 C111 MOG   1        50.057  51.527  28.265  1.00  0.00           C
HETATM   91 H11R MOG   1        49.043  51.946  28.285  1.00  0.00           H
HETATM   92 H11S MOG   1        50.855  52.258  28.159  1.00  0.00           H
HETATM   93 C112 MOG   1        50.330  50.719  29.539  1.00  0.00           C
HETATM   94 H12R MOG   1        49.684  49.839  29.609  1.00  0.00           H
HETATM   95 H12S MOG   1        51.337  50.297  29.430  1.00  0.00           H
HETATM   96 C113 MOG   1        50.260  51.471  30.881  1.00  0.00           C
HETATM   97 H13R MOG   1        49.225  51.464  31.219  1.00  0.00           H
HETATM   98 H13S MOG   1        50.576  52.510  30.739  1.00  0.00           H
HETATM   99 C114 MOG   1        51.158  50.793  31.924  1.00  0.00           C
HETATM  100 H14R MOG   1        50.935  49.725  31.929  1.00  0.00           H
HETATM  101 H14S MOG   1        52.207  51.016  31.677  1.00  0.00           H
HETATM  102 C115 MOG   1        50.870  51.290  33.341  1.00  0.00           C
HETATM  103 H15R MOG   1        49.985  50.811  33.765  1.00  0.00           H
HETATM  104 H15S MOG   1        50.812  52.378  33.386  1.00  0.00           H
HETATM  105 C116 MOG   1        52.088  50.916  34.208  1.00  0.00           C
HETATM  106 H16R MOG   1        52.438  49.903  34.049  1.00  0.00           H
HETATM  107 H16S MOG   1        52.974  51.479  33.901  1.00  0.00           H
HETATM  108 C117 MOG   1        51.866  51.041  35.715  1.00  0.00           C
HETATM  109 H17R MOG   1        51.312  50.125  35.960  1.00  0.00           H
HETATM  110 H17S MOG   1        51.160  51.861  35.878  1.00  0.00           H
HETATM  111 C118 MOG   1        53.164  51.082  36.531  1.00  0.00           C
HETATM  112 H18R MOG   1        53.534  52.109  36.476  1.00  0.00           H
HETATM  113 H18S MOG   1        52.923  50.893  37.579  1.00  0.00           H
HETATM  114 H18T MOG   1        53.875  50.330  36.165  1.00  0.00           H
HETATM  115  O31 MOG   1        49.911  51.117  14.284  1.00  0.00           O
HETATM  116  C31 MOG   1        48.972  50.268  14.725  1.00  0.00           C
HETATM  117  O32 MOG   1        49.120  49.059  14.719  1.00  0.00           O
TER
HETATM  118  C12 OL    1        47.750  50.922  15.323  1.00  0.00           C
HETATM  119  H2R OL    1        47.641  51.885  14.834  1.00  0.00           H
HETATM  120  H2S OL    1        46.918  50.259  15.098  1.00  0.00           H
HETATM  121  C13 OL    1        47.861  51.100  16.844  1.00  0.00           C
HETATM  122  H3R OL    1        48.782  51.568  17.190  1.00  0.00           H
HETATM  123  H3S OL    1        47.980  50.145  17.372  1.00  0.00           H
HETATM  124  C14 OL    1        46.748  51.993  17.395  1.00  0.00           C
HETATM  125  H4R OL    1        46.828  52.991  16.951  1.00  0.00           H
HETATM  126  H4S OL    1        45.763  51.651  17.069  1.00  0.00           H
HETATM  127  C15 OL    1        46.861  52.185  18.915  1.00  0.00           C
HETATM  128  H5R OL    1        46.439  53.131  19.242  1.00  0.00           H
HETATM  129  H5S OL    1        47.923  52.324  19.172  1.00  0.00           H
HETATM  130  C16 OL    1        46.159  51.069  19.700  1.00  0.00           C
HETATM  131  H6R OL    1        45.100  51.117  19.445  1.00  0.00           H
HETATM  132  H6S OL    1        46.652  50.123  19.468  1.00  0.00           H
HETATM  133  C17 OL    1        46.339  51.321  21.203  1.00  0.00           C
HETATM  134  H7R OL    1        46.295  52.405  21.357  1.00  0.00           H
HETATM  135  H7S OL    1        47.369  51.057  21.467  1.00  0.00           H
HETATM  136  C18 OL    1        45.370  50.602  22.163  1.00  0.00           C
HETATM  137  H8R OL    1        44.321  50.900  22.059  1.00  0.00           H
HETATM  138  H8S OL    1        45.541  49.535  22.040  1.00  0.00           H
HETATM  139  C19 OL    1        45.801  50.838  23.588  1.00  0.00           C
HETATM  140  H9R OL    1        46.847  50.757  23.845  1.00  0.00           H
HETATM  141 C110 OL    1        44.927  51.158  24.534  1.00  0.00           C
HETATM  142 H10R OL    1        43.872  51.330  24.344  1.00  0.00           H
HETATM  143 C111 OL    1        45.362  51.371  25.969  1.00  0.00           C
HETATM  144 H11R OL    1        45.519  52.453  26.018  1.00  0.00           H
HETATM  145 H11S OL    1        46.309  50.865  26.148  1.00  0.00           H
HETATM  146 C112 OL    1        44.599  50.869  27.198  1.00  0.00           C
HETATM  147 H12R OL    1        43.569  51.214  27.261  1.00  0.00           H
HETATM  148 H12S OL    1        44.438  49.784  27.209  1.00  0.00           H
HETATM  149 C113 OL    1        45.446  51.269  28.415  1.00  0.00           C
HETATM  150 H13R OL    1        45.515  52.361  28.530  1.00  0.00           H
HETATM  151 H13S OL    1        46.449  50.869  28.315  1.00  0.00           H
HETATM  152 C114 OL    1        44.849  50.603  29.647  1.00  0.00           C
HETATM  153 H14R OL    1        43.763  50.725  29.653  1.00  0.00           H
HETATM  154 H14S OL    1        45.129  49.550  29.605  1.00  0.00           H
HETATM  155 C115 OL    1        45.302  51.225  30.975  1.00  0.00           C
HETATM  156 H15R OL    1        44.846  52.198  31.154  1.00  0.00           H
HETATM  157 H15S OL    1        46.384  51.320  31.002  1.00  0.00           H
HETATM  158 C116 OL    1        45.141  50.292  32.186  1.00  0.00           C
HETATM  159 H16R OL    1        44.088  50.042  32.323  1.00  0.00           H
HETATM  160 H16S OL    1        45.602  49.335  31.926  1.00  0.00           H
HETATM  161 C117 OL    1        45.653  50.783  33.545  1.00  0.00           C
HETATM  162 H17R OL    1        45.371  49.957  34.188  1.00  0.00           H
HETATM  163 H17S OL    1        45.083  51.661  33.865  1.00  0.00           H
HETATM  164 C118 OL    1        47.143  51.131  33.663  1.00  0.00           C
HETATM  165 H18R OL    1        47.504  51.151  34.699  1.00  0.00           H
HETATM  166 H18S OL    1        47.610  50.357  33.044  1.00  0.00           H
HETATM  167 H18T OL    1        47.374  52.114  33.258  1.00  0.00           H
TER
END
```
