## Supplementary Table S3 for "Atomistic Molecular Dynamics Simulations of Trioleoylglycerol – Phospholipid Membrane Systems"

### Experiment 1: TOG 2L System (Conformation Based)

| **Simulation 1 Parameters** | | | | |
| --- | --- | --- | --- | --- |
| **Steps** | **Temperature (K)** | **Ensemble** | **Time (ps)** | **Other Parameters** |
| **Heat1** | **100** | **NVT** | **50** | **dt=0.002,ntt=3,gamma_ln=1.0,ntc=2,ntf=2,tol=0.0000001,iwrap=1** |
| Density | 100 | NVT | 500 |  |
| Equil | 100 | NPT | 1000 | *ntb=2,ntp=1,barostat=2,taup=3.0, |
| PMD | 100 | NPT | 1000 |  |
| **Heat2** | **200** | **NVT** | **100** | **dt=0.002,ntt=3,gamma_ln=1.0,ntc=2,ntf=2,tol=0.0000001,iwrap=1** |
| Density | 200 | NVT | 500 |  |
| Equil | 200 | NPT | 1000 | *ntb=2,ntp=1,barostat=2,taup=3.0, |
| PMD | 200 | NPT | 1000 |  |
| **Heat3** | **298** | **NVT** | **100** | **dt=0.002,ntt=3,gamma_ln=1.0,ntc=2,ntf=2,tol=0.0000001,iwrap=1** |
| Density | 298 | NVT | 500 |  |
| Equil | 298 | NPT | 1000 | *ntb=2,ntp=1,barostat=2,taup=3.0, |
| PMD | 298 | NPT | 1000 |  |
| **ProMD** | 298 | NPT | 210650 |  |
| **Simulation 2 Parameters** | | | | |
| **Steps** | **Temperature (K)** | **Ensemble** | **Time (ps)** | **Other Parameters** |
| **Heat3** | **273** | **NVT** | **100** | **dt=0.002,ntt=3,gamma_ln=1.0,ntc=2,ntf=2,tol=0.0000001,iwrap=1** |
| Density | 273 | NVT | 500 |  |
| Equil | 273 | NPT | 1000 | *ntb=2,ntp=1,barostat=2,taup=3.0, |
| PMD | 273 | NPT | 1000 |  |
| **ProMD** | 273 | NPT | 210750 |  |
| **Simulation 3 Parameters** | | | | |
| **Steps** | **Temperature (K)** | **Ensemble** | **Time (ps)** | **Other Parameters** |
| **Heat4** | **277** | **NVT** | **100** | **dt=0.002,ntt=3,gamma_ln=1.0,ntc=2,ntf=2,tol=0.0000001,iwrap=1** |
| Density | 277 | NVT | 500 |  |
| Equil | 277 | NPT | 1000 | *ntb=2,ntp=1,barostat=2,taup=3.0, |
| PMD | 277 | NPT | 1000 |  |
| **ProMD** | 277 | NPT | 213350 |  |

### Experiment 2: TOG 2L System (Number of TOGs)

| **Simulation Parameters** | | | | |
| --- | --- | --- | --- | --- |
| **Steps** | **Temperature (K)** | **Ensemble** | **Time (ps)** | **Other Parameters** |
| **Heat1** | **100** | **NVT** | **50** | **dt=0.002,ntt=3,gamma_ln=1.0,ntc=2,ntf=2,tol=0.0000001,iwrap=1** |
| Density | 100 | NVT | 500 |  |
| Equil | 100 | NPT | 1000 | *ntb=2,ntp=1,barostat=2,taup=3.0, |
| PMD | 100 | NPT | 1000 |  |
| **Heat2** | **200** | **NVT** | **100** | **dt=0.002,ntt=3,gamma_ln=1.0,ntc=2,ntf=2,tol=0.0000001,iwrap=1** |
| Density | 200 | NVT | 500 |  |
| Equil | 200 | NPT | 1000 | *ntb=2,ntp=1,barostat=2,taup=3.0, |
| PMD | 200 | NPT | 1000 |  |
| **Heat3** | **273** | **NVT** | **100** | **dt=0.002,ntt=3,gamma_ln=1.0,ntc=2,ntf=2,tol=0.0000001,iwrap=1** |
| Density | 273 | NVT | 500 |  |
| Equil | 273 | NPT | 1000 | *ntb=2,ntp=1,barostat=2,taup=3.0, |
| PMD | 273 | NPT | 1000 |  |
| **ProMD** | 273 | NPT | 310750 |  |

### Experiment 3: TOG 2L System (Ensemble Based)

| **Simulation 1 Parameters (NPT Ensemble)** | | | | |
| --- | --- | --- | --- | --- |
| **Steps** | **Temperature (K)** | **Ensemble** | **Time (ps)** | **Other Parameters** |
| **Heat1** | **100** | **NVT** | **50** | **dt=0.002,ntt=3,gamma_ln=1.0,ntc=2,ntf=2,tol=0.0000001,iwrap=1** |
| Density | 100 | NVT | 500 |  |
| Equil | 100 | NPT | 1000 | *ntb=2,ntp=1,barostat=2,taup=3.0, |
| PMD | 100 | NPT | 1000 |  |
| **Heat2** | **200** | **NVT** | **100** | **dt=0.002,ntt=3,gamma_ln=1.0,ntc=2,ntf=2,tol=0.0000001,iwrap=1** |
| Density | 200 | NVT | 500 |  |
| Equil | 200 | NPT | 1000 | *ntb=2,ntp=1,barostat=2,taup=3.0, |
| PMD | 200 | NPT | 1000 |  |
| **Heat3** | **273** | **NVT** | **100** | **dt=0.002,ntt=3,gamma_ln=1.0,ntc=2,ntf=2,tol=0.0000001,iwrap=1** |
| Density | 273 | NVT | 500 |  |
| Equil | 273 | NPT | 1000 | *ntb=2,ntp=1,barostat=2,taup=3.0, |
| PMD | 273 | NPT | 1000 |  |
| **ProMD** | 273 | NPT | 310750 |  |
| **Simulation 2 Parameters (NVT Ensemble)** | | | | |
| **Steps** | **Temperature (K)** | **Ensemble** | **Time (ps)** | **Other Parameters** |
| **Heat1** | **100** | **NVT** | **50** | **dt=0.002,ntt=3,gamma_ln=1.0,ntc=2,ntf=2,tol=0.0000001,iwrap=1** |
| Density | 100 | NVT | 500 |  |
| Equil | 100 | NPT | 1000 | *ntb=2,ntp=1,barostat=2,taup=3.0, |
| PMD | 100 | NVT | 1000 |  |
| **Heat2** | **200** | **NVT** | **100** | **dt=0.002,ntt=3,gamma_ln=1.0,ntc=2,ntf=2,tol=0.0000001,iwrap=1** |
| Density | 200 | NVT | 500 |  |
| Equil | 200 | NPT | 1000 | *ntb=2,ntp=1,barostat=2,taup=3.0, |
| PMD | 200 | NVT | 1000 |  |
| **Heat3** | **273** | **NVT** | **100** | **dt=0.002,ntt=3,gamma_ln=1.0,ntc=2,ntf=2,tol=0.0000001,iwrap=1** |
| Density | 273 | NVT | 500 |  |
| Equil | 273 | NPT | 1000 | *ntb=2,ntp=1,barostat=2,taup=3.0, |
| PMD | 273 | NVT | 1000 |  |
| **ProMD** | 273 | NVT | 310700 |  |

### Experiment 4: Adiposome Mimicking 4L System (Ensemble Based)

| **Simulation 1 Parameters (NPT Ensemble)** | | | | | |
| --- | --- | --- | --- | --- | --- |
| **Steps** | **Temperature (K)** | **Ensemble** | **Time (ps)** | **Restraints** | **Other Parameters** |
| **Heat1** | **100** | **NVT** | **100** | **Yes** | **dt=0.002,ntt=3,gamma_ln=10.0,ntc=2,ntf=2,tol=0.0000001,iwrap=1,** |
| Density1 | 100 | NVT | 1000 | Yes |  |
| Density2 | 100 | NVT | 1000 | Yes |  |
| Density3 | 100 | NVT | 1000 | Yes |  |
| Density4 | 100 | NVT | 1000 | Yes |  |
| Density5 | 100 | NVT | 1000 | Yes |  |
| Equil | 100 | NPT | 1000 | No | *ntb=2,ntp=2,barostat=2,taup=4.0, |
| MD1 | 100 | NPT | 5000 | No |  |
| MD2 | 100 | NPT | 500 | No |  |
| **Heat2** | **200** | **NVT** | **100** | **Yes** | **dt=0.002,ntt=3,gamma_ln=10.0,ntc=2,ntf=2,tol=0.0000001,iwrap=1,** |
| Density1 | 200 | NVT | 1000 | Yes |  |
| Density2 | 200 | NVT | 1000 | Yes |  |
| Density3 | 200 | NVT | 1000 | Yes |  |
| Density4 | 200 | NVT | 1000 | Yes |  |
| Density5 | 200 | NVT | 1000 | Yes |  |
| Equil | 200 | NPT | 1000 | No | *ntb=2,ntp=2,barostat=2,taup=4.0, |
| MD1 | 200 | NPT | 5000 | No |  |
| MD2 | 200 | NPT | 500 | No |  |
| **Heat3** | **273** | **NVT** | **100** | **Yes** | **dt=0.002,ntt=3,gamma_ln=10.0,ntc=2,ntf=2,tol=0.0000001,iwrap=1,** |
| Density1 | 273 | NVT | 1000 | Yes |  |
| Density2 | 273 | NVT | 1000 | Yes |  |
| Density3 | 273 | NVT | 1000 | Yes |  |
| Density4 | 273 | NVT | 1000 | Yes |  |
| Density5 | 273 | NVT | 1000 | Yes |  |
| Equil | 273 | NPT | 1000 | No | *ntb=2,ntp=2,barostat=2,taup=4.0, |
| MD1 | 273 | NPT | 5000 | No |  |
| MD2 | 273 | NPT | 500 | No |  |
| PMD | 273 | NPT | 100900 | No |  |
| **Simulation 2 Parameters (NVT Ensemble)** | | | | | |
| **Steps** | **Temperature (K)** | **Ensemble** | **Time (ps)** | **Restraints** | **Other Parameters** |
| **Heat1** | **100** | **NVT** | **100** | **Yes** | **dt=0.002,ntt=3,gamma_ln=10.0,ntc=2,ntf=2,tol=0.0000001,iwrap=1,** |
| Density1 | 100 | NVT | 1000 | Yes |  |
| Density2 | 100 | NVT | 1000 | Yes |  |
| Density3 | 100 | NVT | 1000 | Yes |  |
| Density4 | 100 | NVT | 1000 | Yes |  |
| Density5 | 100 | NVT | 1000 | Yes |  |
| Equil | 100 | NPT | 1000 | No | *ntb=2,ntp=2,barostat=2,taup=4.0, |
| MD1 | 100 | NPT | 5000 | No |  |
| MD2 | 100 | NPT | 500 | No |  |
| **Heat2** | **200** | **NVT** | **100** | **Yes** | **dt=0.002,ntt=3,gamma_ln=10.0,ntc=2,ntf=2,tol=0.0000001,iwrap=1,** |
| Density1 | 200 | NVT | 1000 | Yes |  |
| Density2 | 200 | NVT | 1000 | Yes |  |
| Density3 | 200 | NVT | 1000 | Yes |  |
| Density4 | 200 | NVT | 1000 | Yes |  |
| Density5 | 200 | NVT | 1000 | Yes |  |
| Equil | 200 | NVT | 1000 | No | *ntb=2,ntp=2,barostat=2,taup=4.0, |
| MD1 | 200 | NVT | 5000 | No |  |
| MD2 | 200 | NVT | 500 | No |  |
| **Heat3** | **273** | **NVT** | **100** | **Yes** | **dt=0.002,ntt=3,gamma_ln=10.0,ntc=2,ntf=2,tol=0.0000001,iwrap=1,** |
| Density1 | 273 | NVT | 1000 | Yes |  |
| Density2 | 273 | NVT | 1000 | Yes |  |
| Density3 | 273 | NVT | 1000 | Yes |  |
| Density4 | 273 | NVT | 1000 | Yes |  |
| Density5 | 273 | NVT | 1000 | Yes |  |
| Equil | 273 | NVT | 1000 | No | *ntb=2,ntp=2,barostat=2,taup=4.0, |
| MD1 | 273 | NVT | 5000 | No |  |
| MD2 | 273 | NVT | 500 | No |  |
| PMD | 273 | NVT | 519100 | No |  |
